# Modulation of fungal virulence through CRZ1 regulated F-BAR-dependent actin remodeling and endocytosis in chickpea infecting phytopathogen *Ascochyta rabiei*

**DOI:** 10.1101/2020.09.17.301176

**Authors:** Manisha Sinha, Ankita Shree, Kunal Singh, Kamal Kumar, Vimlesh Kumar, Praveen Kumar Verma

**Affiliations:** Plant Immunity Laboratory, National Institute of Plant Genome Research, Aruna Asaf Ali Marg, New Delhi-110067, India; Department of Biological Sciences, Indian Institute of Science Education and Research Bhopal (IISER-Bhopal), Bhauri, Bhopal-462066, Madhya Pradesh, India

## Abstract

Polarized hyphal growth of filamentous pathogenic fungi is an essential event for host penetration and colonization. The long-range early endosomal trafficking during the hyphal growth is crucial for nutrient uptake, sensing of host-specific cues, and regulation of effector production. Bin1/Amphiphysin/Rvs167 (BAR) domain-containing proteins mediate fundamental cellular processes, including membrane remodeling and endocytosis. Here, we identified an F-BAR domain protein (ArF-BAR) in the necrotrophic fungus *Ascochyta rabiei* and demonstrate its involvement in endosome-dependent fungal virulence on the host plant, *Cicer arietinum*. We show that ArF-BAR regulates endocytosis at the hyphal tip, localizes to the early endosomes, and is involved in actin dynamics. Functional studies involving gene knockout and complementation experiments reveal that ArF-BAR is essential for virulence. The loss-of-function of ArF-BAR results in delayed formation of first septa from the hyphal tip, crucial for host penetration and proliferation. ArF-BAR was induced in response to oxidative stress and infection and localized to endocytic vesicles within the fungal hyphae. We also show that ArF-BAR is able to tubulate synthetic liposomes, suggesting the functional role of F-BAR domain in membrane tubule formation *in vivo*. Further, our studies identified a stress-induced transcription factor, ArCRZ1 (Calcineurin-responsive zinc finger 1) as key regulator for transcriptional reprogramming of ArF-BAR. We propose a model in which ArCRZ1 functions upstream of ArF-BAR to regulate fungal pathogenesis through a mechanism that involves membrane remodeling and actin cytoskeleton regulation.

**Author summary:** BAR-domain superfamily is known to mold amorphous lipid bilayer into defined tubular shapes and critical for endosome formation and trafficking. Although these processes are studied earlier in the context of their structural and biochemical properties, there is limited evidence on the direct role of F-BAR domain proteins in the pathophysiological development of other economically important fungi. Our study assumes functional significance for plant infection as we identified an F-BAR domain-containing protein that is regulated by a distinct transcriptional network. We characterized F-BAR in a necrotrophic fungal pathogen, *Ascochyta rabiei* that causes the Ascochyta blight (AB) disease in chickpea plants. Additionally, we have also identified a calcium-regulated CRZ1 transcription factor that regulates the transcription of *ArF-BAR*. Our study will help to understand the complex interplay underlying the endosome formation required for fungal virulence.

## Introduction

Polarized hyphal growth is a signature feature of filamentous fungi during host colonization [1]. This feature allows the fungus to sense, coordinate, and respond to an array of cues from the host [2]. Therefore, regulation of hyphal tip growth is one of the major virulence determinants in filamentous fungi. In response to infection, the plant innate immune system recognizes pathogens and fosters effective defense responses. Pathogens must recognize the plant surface cues and counter host-generated defense responses for effective pathogenesis. Additionally, invasive fungi must overcome an intracellular challenge posed by the distance between the elongated invading hyphal architecture and the nucleus [3]. Mounting evidence strongly suggests that the complications associated with this increased distance are overcome by long-distance intracellular communication for rapid and precise transduction of external information [4]. Moreover, the maintenance of extremely polarized hyphal morphology is heavily dependent on endosome trafficking. It is also important to maintain the structural and functional features of a fungal cell [5].

In the case of filamentous fungi, long-distance signaling is mediated by early endosomes (EEs). Besides signal sensing for motor-dependent retrograde signaling, EEs are involved in the recycling of cell wall components, polarisomes and various receptors required for polarized tip growth [6]. Loss of these functions leads to impaired host invasion and virulence [7, 8]. The generation of EEs, which are multipurpose carriers, is a key step in the endocytic pathway and involves the cooperative action of membrane bending and cytoskeleton reorganization. Membrane bending is the cornerstone for the generation of EEs and is regulated by proteins involved in the detection and stabilization of membrane curvature [9, 10].

In animal cells, BAR domain superfamily proteins have been shown to integrate membrane dynamics with cytoskeletal changes [11]. F-BAR domain proteins possess N-terminal α-helical coiled-coil dimers and bind to negatively charged membranes via their positively charged domain. This binding generates membrane curvature and regulates intracellular vesicle trafficking [12, 13]. Depending on the degree of curvature of the dimer, BAR membrane domain superfamily proteins are broadly classified into three families: classical BAR domain, Fer/CIP4 homology BAR (F-BAR) domain and inverse BAR (I-BAR) domain proteins. The N-BAR and F-BAR domains induce positive membrane curvature through concave lipid-binding interfaces and trigger cell membrane invagination. However, I-BAR domains interact with shallow negatively curved membranes through convex lipid-binding interfaces, leading to cell membrane protrusion [14]. The pioneering work in the corn smut fungus, *Ustilago maydis*, revealed the importance of endocytosis for the pathogenic development and virulence of the filamentous fungi by showing impaired early pathogenicity and germination in endocytic mutants [15]. The Cdc15, an F-BAR protein, is involved in cytokinetic ring and septa formation in *U. maydis* [16]. Recent studies in *Magnaporthe oryzae* revealed the importance of N-BAR domain-containing proteins in the growth and virulence of filamentous fungi [17]. Further, the I-BAR protein, Rvs167, was found to be involved in the extension of the rigid penetration peg required during *M. oryzae* invasion [18]. The F-BAR protein like Bzz1p in yeast and Cip4 in *Drosophila a*cts during the early stages of endocytosis in the formation of actin patches [19, 20]. It triggers actin polymerization via the Arp2/3 complex [21]. There has been an intense study on the structural and biochemical properties of BAR domain proteins that contribute to their mode of action. However, there has been limited functional characterization of BAR superfamily proteins in phytopathogenic fungi.

*Ascochyta rabiei* (Pass.) Labr. [teleomorph *Didymella rabiei*], a causal agent of Ascochyta blight (AB) disease in chickpea plants (*Cicer arietinum* L.), is one of the most devastating necrotrophic phytopathogens. *A. rabiei* infects the above-ground parts of this legume plant and greatly reduces the yield of the crop [22]. The fungal hyphae aggregate in the cortical cells of the chickpea plant and differentiate into asexual spores called pycnidia [23, 24]. The genome of *A. rabiei* has been sequenced and analyzed to identify pathogenic determinants [25]. *A. rabiei* has emerged as an interesting model system for elucidating the cell biology, especially the endocytic machinery, during polar growth and pathogenesis in necrotrophic phytopathogenic fungi.

In this study, a F-BAR domain-containing protein, ArF-BAR, was identified in *A. rabiei*. A loss of function mutation of ArF-BAR caused a dramatic reduction in EEs, severely compromised fungal pathogenesis and delayed septa formation. The results showed that the F-BAR domain of the ArF-BAR protein binds to and deforms synthetic liposomes and generates membrane tubules. *ArF-BAR* was induced in response to oxidative stress and infection, and localized to endocytic vesicles within the fungal hyphae. It was also found that *ArF-BAR* expression was regulated by a stress inducible transcription factor, ArCRZ1. The data suggested that ArF-BAR-dependent membrane remodeling combined with actin cytoskeleton dynamics at the fungal hyphal tip was crucial for pathogenesis.

## Results

### *ArF-BAR* expression is induced in response to oxidative stress and infection

Transcriptome analysis of *A. rabiei* during oxidative stress has provided a greater understanding of the survival strategies used by necrotrophic fungi against host-generated oxidative stress [26]. The study by Singh et al. [26] revealed that 70 unigenes were upregulated under oxidative stress conditions. Of these 70 unigenes, a clone resembling “ST47_g8005” of the sequenced *A. rabiei* genome [27], showed early upregulation against oxidative stress. The deduced amino acid sequence of this gene revealed a protein with four distinct domains: an N-terminal F-BAR domain, a unique protein kinase C1 domain, and two consecutive C-terminal SH3 domains (Fig 1A). Henceforth, this protein has been named as ArF-BAR. Phylogenetic analysis of selected pathogenic fungi and other eukaryotes revealed that ArF-BAR shared sequence identity with proteins of many closely related phytopathogens (S1 Fig). ArF-BAR was found to share approximately 33% and 41% sequence identity with *Saccharomyces cerevisae* BZZ1p and *Drosophila melanogaster* Cdc42-Interacting Protein 4 (CIP4) proteins, respectively (S2 Fig).

**Fig 1.**
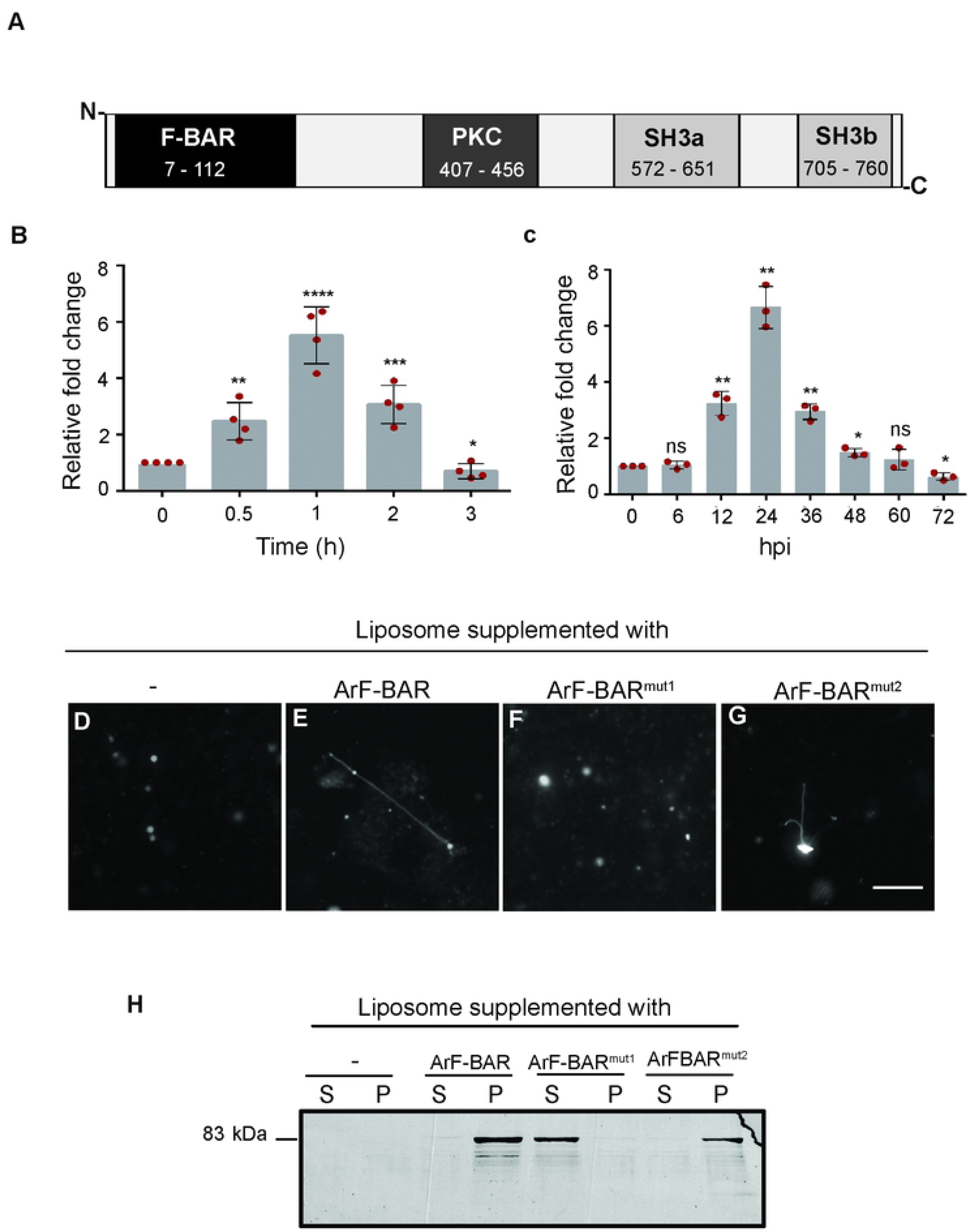
ArF-BAR is a bonafide stress induced membrane tubulating protein. (A) Schematic representation of ArF-BAR full-length protein domains. (B) Relative fold change in the transcript level of Ar*F-BAR* under menadione treated condition as analysed with qRT-PCR. (C) Relative fold change in the transcript level of Ar*F-BAR* during *in planta* infection as analysed with qRT-PCR. (D-G) Membrane deforming activity of ArF-BAR. The bar represents 5 mm. (D) In the absence of recombinant protein spherical liposome are found, (E,G) Synthetic liposomes deforms into tubules with of addition of ArF-BAR and ArF-BAR^mut2^ recombinant proteins, (F) Liposome tubulation activity is lost in ArF-BAR^mut1^. (G) Liposome co-sedimentation assay, where the ArF-BAR and ArF-BAR^mut2^ recombinant protein bound with lipid membranes (liposome) were found in pellet, while membrane binding activity is lost in ArF-BAR^mut1^ protein. Statistical analysis was performed using Student’s t-test one tailed compared with its control. Significant differences are indicated as \**p* < 0.03; \*\**p* < 0.003; \*\*\**p* < 0.0005; \*\*\*\**p* < 0.0001; ns = non-significant. Data is the mean of three independent biological replicates with error bars ± representing standard deviation. Red dots represents the value of each biological replicate/ sample used for the quantitative analysis.

To validate the results of the transcript profiling, quantitative real-time PCR (qRT-PCR) was performed using primers specific to the F-BAR domain encoding region. The qRT-PCR experiments revealed a higher expression level of the *ArF-BAR* transcript after a 1 h treatment of menadione, which is an oxidative stress generator (Fig 1B). To directly assess *ArF-BAR* transcript induction during pathogenesis in a susceptible chickpea variety (PUSA-362), the time course of *ArF-BAR* transcript expression was measured following plant inoculation with conidial suspension using qRT-PCR. Consistent with the previous observations, a significant increase in the *ArF-BAR* transcript level was found following infection. The maximum transcript level was at 24 hours post-infection (hpi; Fig 1C), which is within the critical time period for spores to germinate on the host surface [27].

### ArF-BAR is a membrane tubulating protein

To understand whether ArF-BAR has a functional F-BAR domain, synthetic liposomes containing Rhodamine B-conjugated PE were incubated with purified His-tagged recombinant ArF-BAR protein (S3A Fig). The spherical liposomes transformed into an intense network of narrow tubules within 30 min of incubation (Fig 1D and 1E). This indicated that ArF-BAR was capable of robust liposome tubulation activity (S3B Fig). Additionally, the sequence alignment of the ArF-BAR, F-BAR domain with that of other F-BAR domain-containing proteins, revealed the presence of positively charged conserved residues (S2 Fig). Earlier, the positively charged present in the F-BAR domains of CIP4 and Syndapin, have been shown to interact with negatively charged phospholipids to induce membrane curvature [28, 29]. Since ArF-BAR has conserved positively charged amino acids at positions 57, 58, 131 and 132 (S2 Fig), the role of these residues in membrane deformation were evaluated. This evaluation was performed by replacing the residues with glutamate (K57E, K58E, R131E and K132E) via site-directed mutagenesis, and the mutant was referred to as ArF*-*BAR^mut1^ (S3C Fig). Predictably, it was found that substituting the conserved lysine/arginine with glutamate abolished the tubulation activity of the F-BAR domain (Fig 1F). Protein kinase C (PKC) proteins are diversely known to interact with diacylglycerol (DAG), which has a role in membrane interactions. To abolish the involvement of the unique protein kinase C1 domain of ArF-BAR in membrane deformation, the W428 and L430 residues were replaced with G428 and G430, respectively [30], and the mutant form was referred to as ArF-BAR^mut2^ (S3 D). Further, this mutated protein was used to assess the liposome tubulation activity, and a similar activity to that of the native protein was found (Fig 1G). To test whether the tubulation activity of ArF-BAR correlated with its lipid binding ability, a liposome co-sedimentation assay was used (Fig 1H). Compared to the native ArF-BAR protein, ArF-BAR^mut1^ showed reduced lipid sedimentation efficiency. In contrast, the sedimentation was unaffected with ArF-BAR^mut2^. This indicated the importance of the direct interaction of the BAR domain with the lipids in liposome tubulation (Fig 1H). Together, these results suggest that the F-BAR domain of ArF-BAR, along with its positively charged residues, plays a major role in interacting with the lipid membrane for binding and deformation.

### ArF-BAR is required for the virulence of *A. rabiei* in chickpea

To elucidate the biological importance of ArF-BAR in *A. rabiei* pathogenesis, an *A. rabiei* knockout mutant that lacked the entire *ArF-BAR* gene was generated (*Δarf-bar*). The *ArF-BAR* gene was targeted for deletion using a homologous recombination approach. The results confirmed that the open reading frame (ORF) of *ArF-BAR* had been successfully replaced with a single copy of the hygromycin resistance gene (*Hph*, S4A and S4B Fig). Simultaneously, *Δarf-bar* mutant strain was complemented with a T-DNA cassette that expressed *ArF-BAR* under its own promoter (*Δarf-bar/ArF-BAR*, S4B and S5A Fig).

Notably, the knockout mutants showed reduced radial growth compared to wild-type *A. rabiei* (WT) and this radial growth was restored in the *Δarf-bar/ArF-BAR* complementation mutant (Fig 2A). To determine the virulence of the fungal strains, an *in planta* infection bioassay was performed on a susceptible chickpea variety. Typical AB disease symptoms were observed on plants challenged with the *A. rabiei* (WT) and *Δarf-bar/ArF-BAR,* but not on plants challenged with *Δarf-bar* (Fig 2B). The degree of pathogenicity was measured according to lesion number and lesion size, which were compared among the *A. rabiei* (WT) and mutants at 144 hpi. The lesion number per plant in *Δarf-bar* was significantly lower than in the *A. rabiei* (WT, Fig 2C). The mean lesion size was also much lower in *Δarf-bar* than in the *A. rabiei* (WT, Fig 2d). However, for the *Δarf-bar/ArF-BAR* complementation mutant, the disease symptoms were comparable to those of the *A. rabiei* (WT). Overall, the disruption of the *ArF-BAR* gene significantly impaired the pathogenicity of *A. rabiei*.

**Fig 2.**
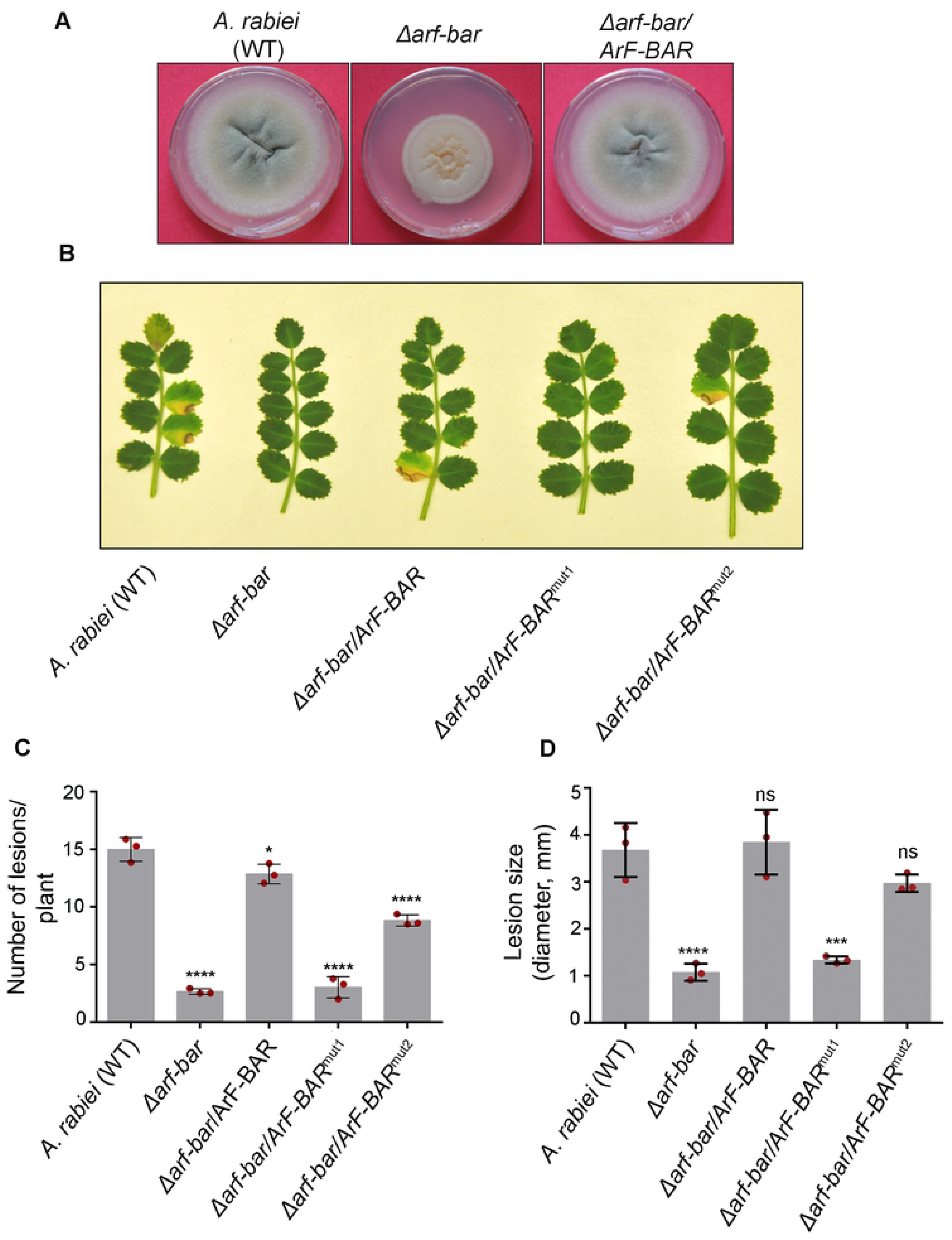
ArF-BAR is required for pathogenicity of *A. rabiei* in chickpea. (A) *A. rabiei* (WT), *Δarf-bar* and *Δarf-bar/ArF-BAR* mutant strains were grown on PDA plate for 7 days at 22°C. (B) Representative image of the AB disease symptoms on susceptible chickpea plants at 6 dpi, obtained after inoculation with *A. rabiei* (WT) and mutant strains conidia. Loss in pathogenicity was observed in *Δarf-bar and Δarf-bar/ArF-BAR*^mut1^. (C) The number of lesions per plant were counted when treated with *A. rabiei* (WT), mutant, and mutant complemented strains conidia. The graph represents the mean and standard deviation of three independent biological replicates, counting at least 10 plants in each replicate. (The bar graph corresponds to-14.99 ± 0.84 in WT; 2.66 ± 0.19 in *Δarf-bar*; 12.86 ± 0.69 in *Δarf-bar/ArF-BAR*; 3.02 ± 0.74 in *Δarf-bar/ArF-BAR*^mut1^and 8.833 ± 0.04 in *Δarf-bar/ArF-BAR*^mut2^, respectively). (D) Infection size was calculated with lesion diameter (The bar in graphs corresponds to, WT = 3.67 mm ± 0.46; *Δarf-bar* = 1.07 mm ± 0.14; *Δarf-bar/ArF-BAR* = 3.84 ± 0.54; *Δarf-bar/ArF-BAR*^mut1^ = 1.33 ± 0.06 and *Δarf-bar/ArF-BAR*^mut2^ = 2.97 ± 0.15). Statistical analysis was performed using ordinary one-way ANOVA compared with its control (\*\*\*\**p* < 0.0001; \**p* < 0.05). Data is the mean of three independent biological replicates with error bars ± representing standard deviation. Red dots represents the value of each biological replicate/ sample used for the quantitative analysis.

To ascertain the role of the evolutionary conservation of the F-BAR and unique PKC domains in contributing to fungal pathogenicity, the two domains were independently inactivated by site-directed mutagenesis as described earlier. The *Δarf-bar* knockouts were complemented with mutated *ArF*-*BAR*s encoding for mutated F-BAR (*Δarf-bar/ArF-BAR*^mut1^) and PKC domains (*Δarf-bar/ArF-BAR*^mut2^; S4B, S5B and S5C Fig). The degree of pathogenicity in *Δarf-bar/ArF-BAR*^mut1^ strain was lower than that of the WT. Meanwhile, in the case of *Δarf-bar/ArF-BAR*^mut2^, the number of lesions per plant was significantly lower than in the WT, but there was no significant difference in the size of the lesions (Fig 2B, 2C and 2D). The severity of the disease symptoms increased in the plants after 10 days post infection (dpi), but no significant differences were observed for plants challenged with *Δarf-bar* (S6 Fig). Together, these results confirm that *ArF-BAR* is an important pathogenicity determinant and that the F-BAR domain is indispensable for fungal pathogenesis.

It was subsequently hypothesized that the loss in pathogenicity of *Δarf-bar* could have resulted from at least two factors: a) knockout mutants of *ArF-BAR* may have had a compromised ability to penetrate host tissue, or b) the virulence phenotype of the mutant was a consequence of a reduction in fungal viability. To address these possibilities, the depth of hyphal penetration in chickpea leaves infected with the WT and *Δarf-bar* mutant strains were examined and compared. The leaves of the chickpea plants challenged with the fungi were subjected to wheat germ agglutinin (WGA-488) staining. At 48 hpi, the infected leaves were stained with WGA-488 to enable visualization of the fungus [30]. The infected leaves were optically sectioned using confocal microscopy starting from the surface of the leaves. It was observed that hyphae of the wild-type strain efficiently penetrated into the chickpea leaves up to depths of 20.25 µm ± 3.92 (mean ± SD; *n* = 3). In stark contrast, the hyphae of the *Δarf-bar* mutant were unable to penetrate beyond 9.83 µm ± 2.54 (mean ± SD; *n* = 3, S7A and S7C Fig). Thus, the depth and efficiency of hyphal penetration by *Δarf-bar* were significantly impaired (S7B and S7C Fig).

Further, to investigate the direct involvement of *ArF-BAR* in fungal viability under oxidative stress conditions, radial growth assays were performed for WT, *Δarf-bar*, *Δarf-bar/ArF-BAR*, *Δarf-bar/ArF-BAR*^mut1^ and *Δarf-bar/ArF-BAR*^mut2^ strains. These strains were inoculated on either potato dextrose agar (PDA) or PDA supplemented with menadione (250 µM and 500 µM) or H_2_O_2_ (2 mM). The diameter of radial growth was analyzed at 10 dpi. The mycelial growth of *Δarf-bar* was considerably reduced compared to that of the WT. The mycelial growth was restored in the *Δarf-bar/ArF-BAR* and *Δarf-bar/ArF-BAR*^mut2^ mutants. However, the mycelial growth of *Δarf-bar/ArF-BAR*^mut1^ was unable to match that of the WT. Exposure to oxidative stress led to a greater growth inhibition rate in *Δarf-bar* than in the WT (S8A and S8B Fig). The involvement of *ArF-BAR* in oxidative stress tolerance was confirmed by the complementation mutant, *Δarf-bar/ArF-BAR,* which showed similar resistance to the WT (S8A and S8B Fig). Further, exposure to oxidative stress led to greater growth inhibition in *Δarf-bar/ArF-BAR*^mut1^ than in the WT. However, the growth inhibition of *Δarf-bar/ArF-BAR*^mut2^ was similar to that of WT. Consistent with the previous findings, these results strongly support the hypothesis that *ArF-BAR* is a positive regulator of pathogenicity in *A. rabiei* and is required for the viability of the fungus under stress conditions.

### Absence of *ArF-BAR* delays septa formation

The next aim was to determine the subcellular localization of ArF-BAR. In this regard, an enhanced yellow fluorescent protein (EYFP) was used to create the fusion protein, ArF-BAR::EYFP, which was transiently expressed in *Δarf-bar*. The bioimaging of the fluorescently tagged ArF-BAR showed punctate distribution throughout the cytoplasm of the fungal hyphae. The chimeric protein was mostly concentrated at the growing hyphal tip and at the septa (Fig 3A and 3B). Fungal hyphae transformed with chimeric ArF-BAR^mut1^::EYFP exhibited disrupted localization of these punctate structures and the fluorescence was completely diffused throughout the cytoplasm. However, the distribution of the fluorescent puncta was unaffected in hyphae transiently expressing ArF-BAR^mut2^::EYFP. The punctate structures were distributed throughout the cytoplasm and were prominently concentrated at the growing hyphal tip and septa (Fig 3A). Further, to validate the spatiotemporal distribution pattern of the ArF-BAR protein during infection, fungal spores expressing ArF-BAR::EYFP were allowed to infect susceptible chickpea leaves. Microscopic observation of these hyphae infected chickpea leaves revealed a distribution pattern that was similar to that of the ArF-BAR::EYFP expressing hyphae growing on glass slides; the chimeric protein was predominantly distributed at the hyphal tip and septa (S9 Fig).

**Fig 3.**
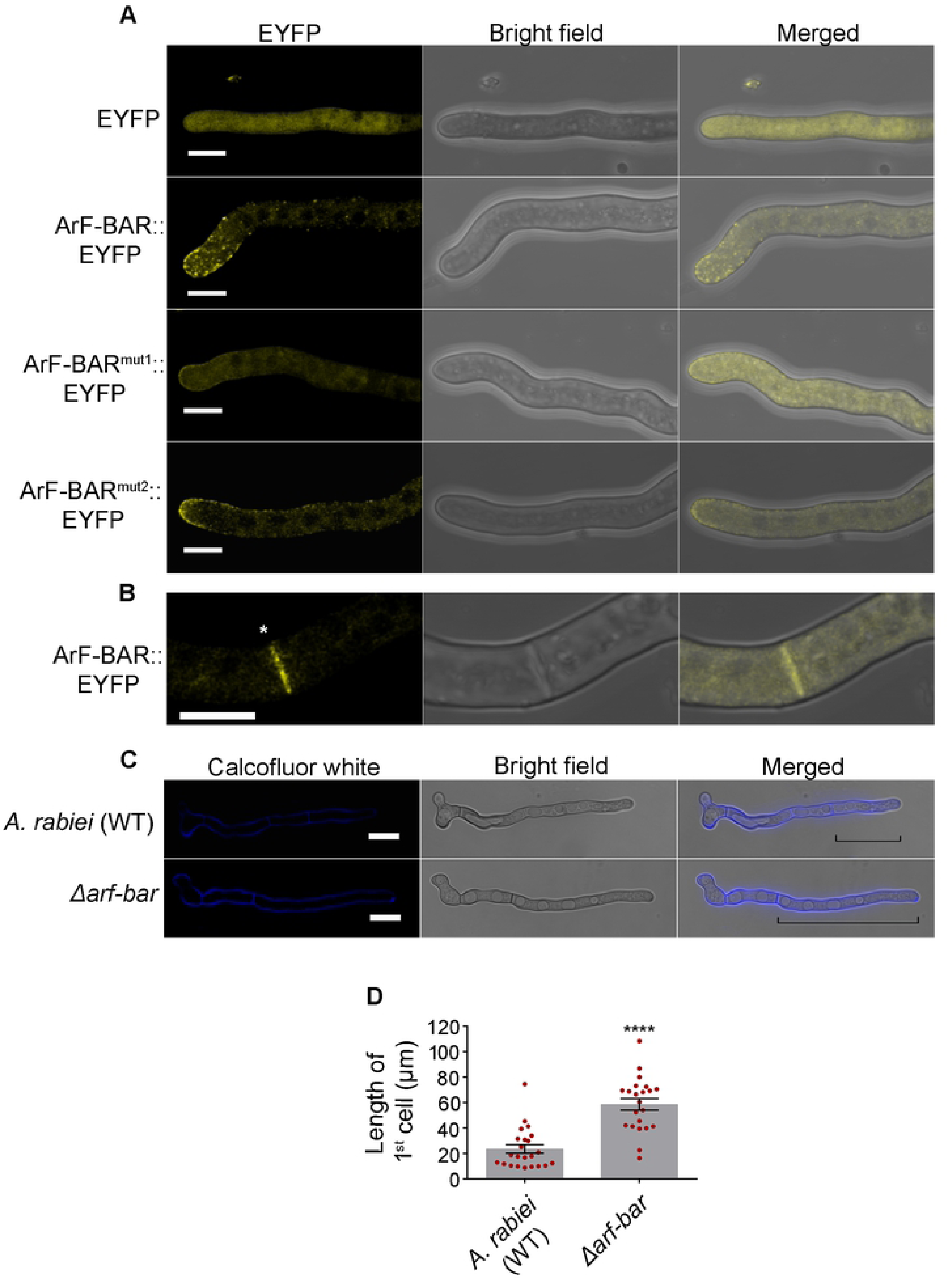
*Δarf-bar* perturbs septa formation. (A) Ectopically expressed ArF-BAR::EYFP and ArF-BAR^mut2^::EYFP chimeric proteins are distributed at the hyphal tip. Hyphae expressing ArF-BAR^mut1^::EYFP, was diffused throughout the fungal hyphae. The representative images were captured at 12 h post germination of fungal conidia on microscopic coverslip. Scale bar represents 5 µm. (B) ArF-BAR::EYFP protein is uniformly distributed at fungal septum, where the scale bar = 5 µm. Star represents the septa. (C) Calcofluor white stained *A. rabiei* (WT) and *Δarf-bar* hyphae after 12 h post germination. Increase in the distance of first septa from the polarised end was found in *Δarf-bar.* Line marks the position of septa in both *A. rabiei* (WT) and *Δarf-bar.* Scale bar = 10 µm. (D) Bar graph represents the mean with SEM, of length of the first septa formed from the growing end. Significance of the difference in length of first cell from the growing tip region compared to WT was calculated using one-tailed paired t-test, (\*\*\*\**p* < 0.001). Red dots represents the value of each biological replicate/ sample used for the quantitative analysis.

The arrangement of septa within the fungal hyphae of both the WT and the *Δarf-bar* strain were examined. Microscopic analysis of the *Δarf-bar* mutant using calcofluor white, which precisely stains cell wall components, showed that the filaments lacked regularly spaced septa. Interestingly, the distance of the first septum from the growing hyphal tip (polarized end) was significantly greater in the *Δarf-bar* mutant (58.5µm ± 4.65; mean ± SEM) than in the WT (23.5µm ± 3.23; mean ± SEM; Fig 3C and3D). Overall, these results reveal that the *ArF-BAR* gene is necessary for appropriate fungal architecture, which is, in turn, important for host penetration and virulence.

### ArF-BAR mediates early endosome biogenesis and endocytosis

Since F-BAR domain proteins are known to form a canonical banana-shaped fold and to dimerize [31], the dimerization of ArF-BAR was confirmed using a yeast two-hybrid (Y2H) system. This evidence indicated the evolutionarily conserved nature of the BAR protein function (S10A Fig). To further gain insights of ArF-BAR in endocytosis an endocytic tracer dye, N-(3-triethylammoniumpropyl)-4-(p-diethyl-aminophenyl-hexatrienyl)pyridiniumdi bromide (FM4-64), was used [32]. Both WT and *Δarf-bar* were stained, and the microscopic analysis revealed visible internal staining of the hyphae. The results suggested that in WT-hyphae, the dye was rapidly internalized. In contrast, in *Δarf-bar* strain, no such obvious internalization was observed, even after 10–15 min of FM4-64 incubation (Fig 4A). To quantify these findings, the mean fluorescence intensity was determined (Fig 4A). The results provided evidence for the involvement of *ArF-BAR* in the endocytic mechanism.

**Fig 4.**
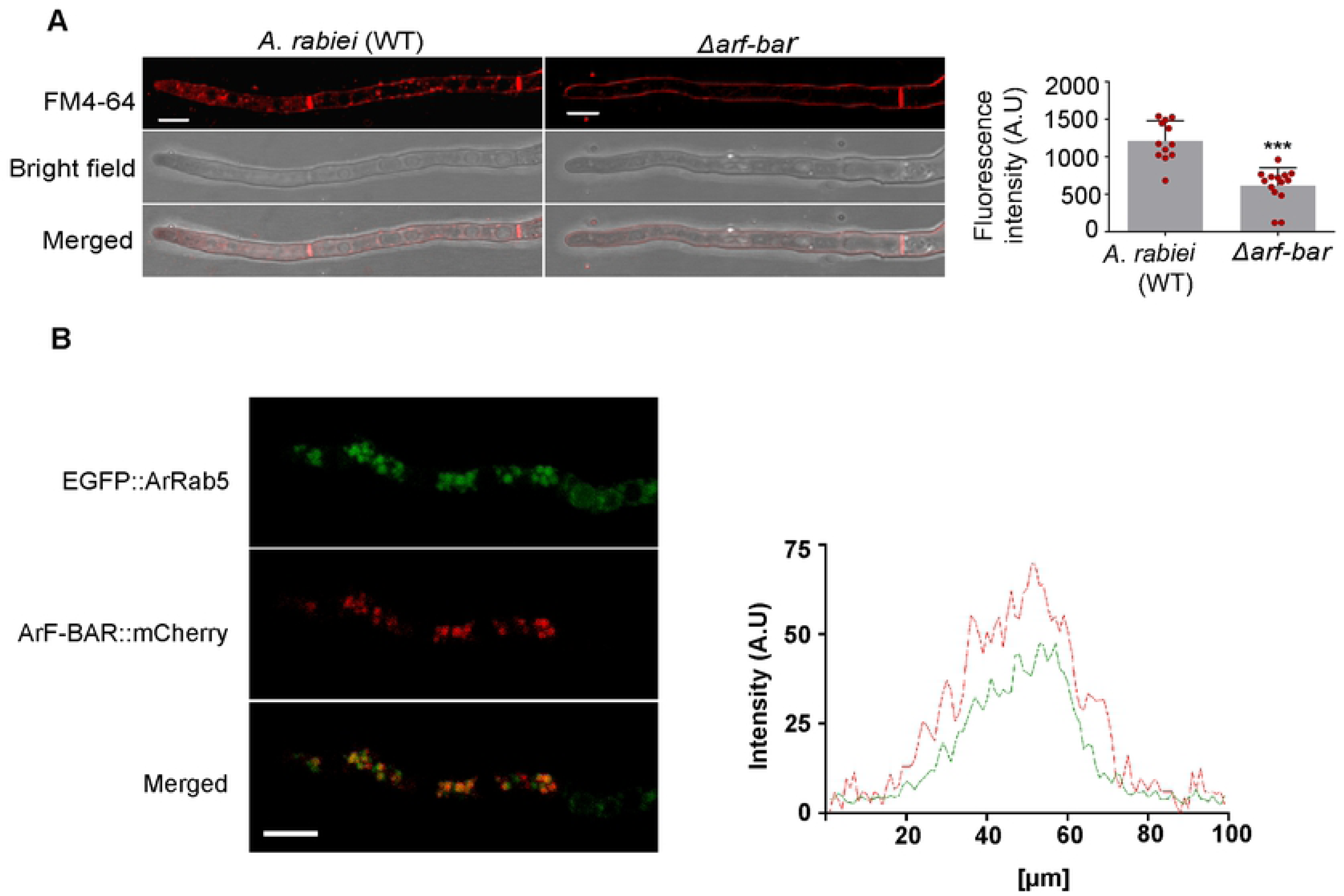
ArF-BAR is facilitates early endosomal biogenesis and endocytosis. (A) Confocal images of *A. rabiei* (WT) and *Δarf-bar* hyphae after 10 min incubation with FM4-64 to acquire internalization capacity. Scale bar = 10 µm. Right panel represents the difference in fluorescence intensity of FM4-64 in *A. rabiei* (WT) and *Δarf-bar* hyphae after internalization. The mean fluorescence intensity was quantified using one-tailed paired t-test, \*\*\**p* < 0.0002. (B) Confocal images showing co-localization of ArF-BAR::mCherry with EGFP::ArRab5 (n = 22), scale bar = 5 µm. Right panel shows the positive correlation between the fluorescence intensity of ArF-BAR (Red) with ArRAB5 (Green).

The central components of the endocytic pathway are the EEs, where the small GTPase Rab5 plays a major regulatory role in biogenesis [33]. *ArRab5,* an orthologue of *Rab5,* was identified in the *A. rabiei* through NCBI blast search, against *Rab5* of *M. oryzae*, and *U. maydis*. The relationship between ArF-BAR-associated endocytosis and ArRab5-associated early endosomes was determined using a double-labeling experiment. *ArF-BAR* was tagged with *mCherry* and *ArRab5* with *EGFP.* This was followed by sequential transformation into the WT. The coalescence of fluorescence obtained showed perfect positive correlation between the two fusion proteins (Fig 4B).

Early endosomes mature to late endosomes followed by the replacement of Rab5 to Rab7 [34]. Thus, we aimed to determine whether the punctate distribution of ArF-BAR was associated with all endocytic vesicles or specifically to the EEs. In this context, similar to Rab5 orthologue, an orthologue of *Rab7* was identified in *A. rabiei*, and was tagged with EYFP (*ArRab7:EYFP*). A similar double-labeling experiment was performed that showed that least correlation between the ArF-BAR::mCherry and ArRab7::EYFP fusion proteins (S10B Fig). In summary, these results uncover a novel role of ArF-BAR proteins; they specifically bind to early endocytic vesicles, regulate the biogenesis of EEs, and mediate their motility during endocytosis.

### ArF-BAR modulates the actin cytoskeleton

The presence of SH3 (SRC homology 3) domains in F-BAR proteins is well documented for their relationship with the actin cytoskeleton via interactions with the Arp2/3 complex activator Wiskott-Aldrich syndrome protein (WASp) [35]. Since ArF-BAR in *A. rabiei*, contains two consecutive SH3 domain at its C-terminus, we initially hypothesized for the probable existence of interactions of ArF-BAR with that of ArActin. Using Y2H system, it was shown that ArF-BAR does not interact directly with actin (Fig 5A), rather it physically interacts with WASp through its SH3 domain (572-760 amino acids; Fig 5B, S11A and S11B Fig). Further, to check the role of ArF-BAR in actin polymerization, a well-established *in vitro* actin polymerization assay was performed. The kinetics of actin polymerization was monitored by the increase in the fluorescence of pyrene-labeled actin. The effect of purified ArF-BAR protein on actin nucleation (actin, Arp2/3 and WASp) was tested using a minimal set of components for all reactions. Interestingly, the addition of purified recombinant protein led to an increase in the actin polymerization rate (Fig 5C). By increasing the concentration of purified protein, a significant gradual activation in the rate of actin polymerization was observed (Fig 5C). These results strongly suggest that ArF-BAR plays an active role in WASp-dependent actin polymerization.

**Fig 5.**
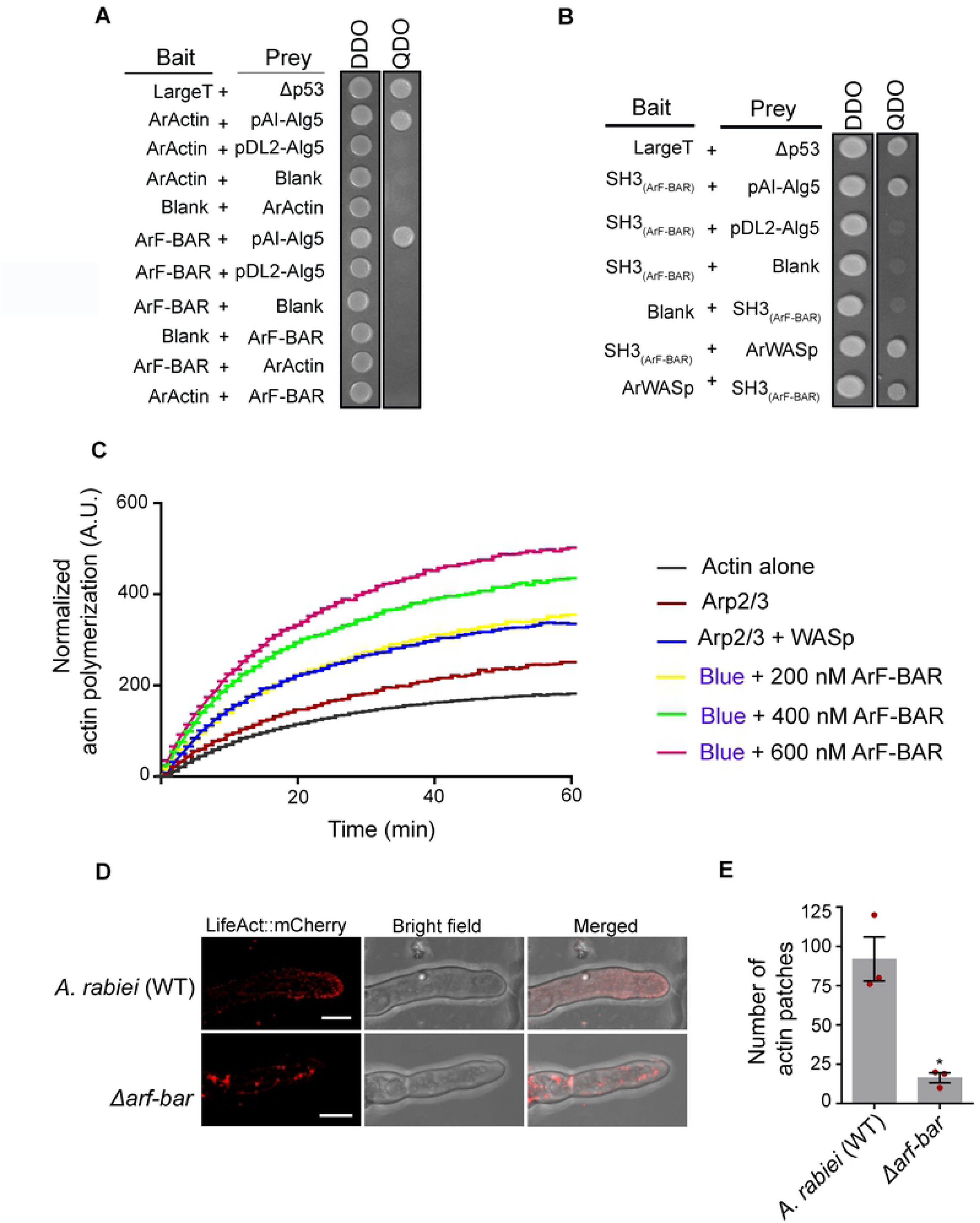
ArF-BAR modulates the actin cytoskeleton. (A) Yeast two-hybrid showing no physical interaction of ArF-BAR with ArActin, as the colonies growing on DDO (SD/-L/-W) fail to grow on QDO (SD/-L/-W/-A/-H) (B) ArF-BAR exhibits positive interaction with ArWASp through its SH3 domain [SH3_(ArF-BAR)_; 508-760 amino acids]. The representative images were photographed 48 h after yeast clones spotting. These results are confirmed with three independent biological replicates. (C) Kinetics of WASp and Arp2/3 mediated actin polymerization of ArF-BAR, measured with the change in fluorescence of pyrene-actin. For all *in vitro* actin polymerization assays; 4 µM pyrene labelled actin, 13 nM Arp2/3 and 15 nM WASp were used. All the experiments was performed in three replicates. (D) Fluorescence image of LifeAct::mCherry expressing in fungal hyphae WT and *Δarf-bar*, where discrete actin patches and cables are visible in WT however actin patches are sparsely visible in *Δarf-bar*. Scale bar = 5 µm (n = 10).

Appropriate organization of actin is required for vesicular dynamics, organelle movement and cytokinesis. Actin microfilaments or F-actin are organized into higher order structures comprising of actin patches, cables and rings that serve as the track for long distance transport [36]. To elucidate the relative importance of ArF-Bar in actin organization the actin dynamics were compared in the WT and *Δarf-bar* mutants. Here, we took advantage of LifeAct, an actin binding peptide fused with a fluorescence protein. LifeAct has been successfully employed for *in vivo* visualization of actin filaments and dynamics in a variety of organisms including fungi and plants [37]. In this study, a *LifeAct:mCherry* fusion construct was generated and transformed into both the WT and *Δarf-bar* mutants to visualize cytoplasmic actin in the fungal hyphae. Confocal microscopy revealed discrete actin patches, and cables in the WT hyphae (Fig 5D). In contrast, the actin patches were rarely visible in the *Δarf-bar* mutant (Fig 5D and 5E). Moreover, the actin cables were dramatically disorganized in the *Δarf-bar* mutant. Although ArF-BAR does not directly interact with actin, it regulates actin polymerization and the actin cytoskeleton through its association with WASp in the growing fungal hyphae.

### ArCRZ1 is a potential transcriptional regulator of *ArF-BAR*

Thus far, the findings of the present study have revealed the importance of *ArF-BAR* in the regulation of endocytic pathways, which is crucial for the pathogenesis of *A. rabiei*. However, the transcriptional regulatory machinery associated with the endocytic pathway in filamentous pathogenic fungi is largely unknown. Hence, the transcriptional mechanism associated with the regulation of *ArF-BAR* in response to infection was subsequently analyzed. Binding motifs for fungal transcription factors (TFs) were identified in the upstream regulatory sequences of *ArF-BAR*. Seven different putative TF binding sites were identified (S1 Table). Among these, three binding sites, positioned at -106, -136, and -254 upstream of the *ArF-BAR* promoter, were found for calcineurin-responsive zinc finger transcription factor 1 (CRZ1; Fig 6A).

**Fig 6.**
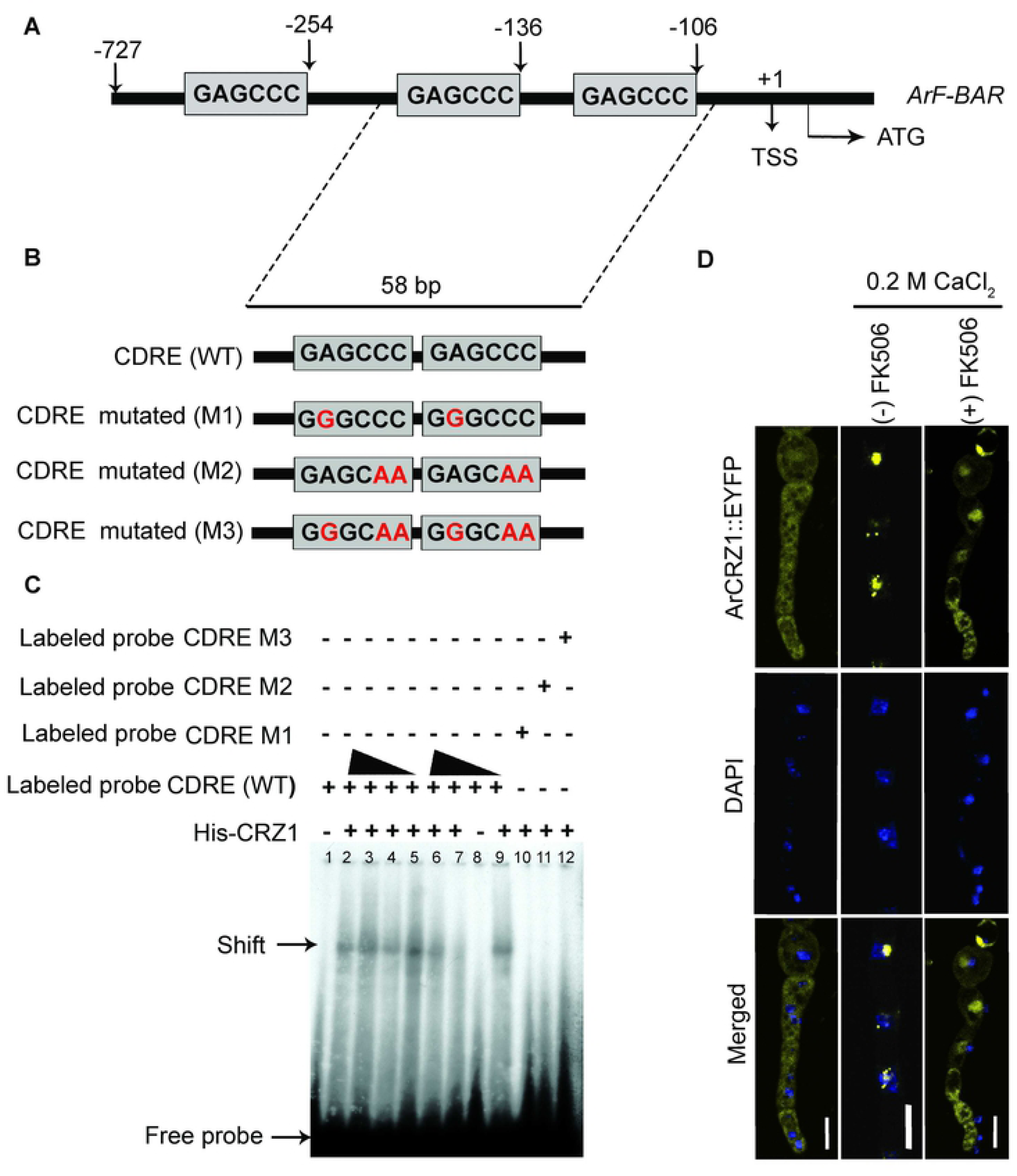
ArCRZ1 functions in the upstream of *ArF-BAR*. (A) Schematic representation of ArCRZ1 TF binding sites at the 5′ regulatory region of *ArF-BAR* gene. (B) Schematic representation of 58 bp WT and mutated Calcium Dependent Response Element (CDRE) probes derived from 5′ regulatory region of *ArF-BAR* gene. (C) The electrophoretic mobility shift assay (EMSA) of recombinant ArCRZ1 with WT or mutated CDRE probes showing the specific binding specificity of ArCRZ1 CDREs. Mutated sites are depicted in red colour. His-purified ArCRZ1 recombinant protein in 500, 600 and 200 ng was used in lane 2, 3, 4, respectively. Same protein in 800, 200 and 100 ng amount was used in lane 5, 6, and 7, respectively. The lane 1 and 8 contains only probe. The lane 9 has WT CDRE, and lane 10, 11 and 12 have mutated CDRE. The plus (+) and minus sign (-) represents the presence and absence of proteins. This binding experiment was replicated in triplicates. (D) Confocal images showing the cytosolic distribution of ArCRZ1::EYFP in the absence of stress condition and nuclear localization of ArCRZ1::EYFP in hyphae under stress condition (0.2 M CaCl_2_). Overnight grown fungal hyphae was exposed to 0.2 M CaCl_2_ two minutes prior to the microscopy. DAPI fluorescence was simultaneously recorded. (Scale bar = 5 µm). Right panel in the confocal images shows the cytosolic distribution of ArCRZ1::EYFP under stress condition (200 mM CaCl_2_) with calcineurin inhibition in the presence of FK506 (5 µg/µl). Fungal hyphae was exposed to FK506 for 5 min, prior to microscopy.

CRZ1 is an evolutionarily conserved TF from yeast to mammals. CRZ1 was chosen for analysis because it regulates the expression of various genes involved in stress tolerance [38] and is widely known to translocate inside the nucleus with an increase in cytosolic Ca^2+^ ion concentration. CRZ1 of *A. rabiei* (*ArCRZ1*; ST47_g3738) possesses a serine-rich region (SRR), two consecutive calcineurin docking domains (CDD), characterized by PxlxlT motif (PRlLPQ and PElNlD) and a single C_2_H_2_ zinc finger motif (S12A Fig). To determine the role of ArCRZ1 in the transcriptional regulation of *ArF-BAR*, the binding of ArCRZ1 to the regulatory sequences of *ArF-BAR* was confirmed. This confirmation was performed via an electrophoretic mobility shift assay (EMSA) using recombinant His-tagged ArCRZ1 proteins. Shifting was observed for the DNA fragment possessing the calcineurin-dependent response element (CDRE) in the presence of purified His-ArCRZ1. However, mutations in this CDRE resulted in the abolishment of binding (Fig 6B and 6C).

Subsequently, the sub-cellular localization of ArCRZ1 under Ca^2+^ and oxidative stress conditions was determined. The WT was transformed with a translational fusion of *ArCRZ1* with *EYFP* towards C-terminus. Confocal microscopy revealed the elegantly concentrated nuclear localization of the ArCRZ1::EYFP signal under stress conditions (Fig 6D). Interestingly, the ArCRZ1::EYFP fusion protein was uniformly distributed within the cytoplasm in the absence of these stresses (Fig 6D). Since, the nuclear translocation of ArCRZ1 is mediated by a phosphatase (calcineurin), a chemical genetics approach was used to confirm the relationship between calcineurin and ArCRZ1. Here, the immunosuppressant FK506, which is a potent calcineurin inhibitor [39], was used to silence the enzymatic activity of calcineurin. FK506 potentially inhibited nuclear translocation of ArCRZ1::EYFP, which indicates that calcineurin plays a role in the nuclear translocation of ArCRZ1 under stress conditions (Fig 6D).

To corroborate and extend these findings during infection, susceptible plants were challenged with fungal conidia expressing ArCRZ1::EYFP, and nuclear localization of EYFP was observed (S12B and S12C Fig). Overall, these results confirm the evolutionarily conserved signaling of calcineurin-dependent ArCRZ1 under oxidative stress conditions.

To uncover the functional regulation of the *ArF-BAR* gene mediated by *ArCRZ1*, a targeted deletion of the *ArCRZ1* gene was generated (*Δarcrz1*), followed by complementation with*ArCRZ1* controlled by its own promoter (*Δarcrz1/ArCRZ1*; S12C-S12E Fig). Interestingly, no significant difference in radial growth diameter was observed between *Δarcrz1* and the WT (Fig 7A). The expression pattern of *ArF-BAR* in the WT and *Δarcrz1* was analyzed using semi-quantitative RT-PCR. The results clearly showed significant reduction in*ArF-BAR* gene expression in the *Δarcrz1* mutant. Imposing oxidative stress to both the WT and the *Δarcrz1* mutant via menadione treatment revealed an upregulation of the *ArF-BAR* transcript in the WT. This upregulation was found to be completely abolished in the *Δarcrz1* mutant (Fig 7B). Overall, these results substantiate that the novel transcriptional regulation of *ArF-BAR* expression is mediated by the TF ArCRZ1 under oxidative stress conditions.

**Fig 7.**
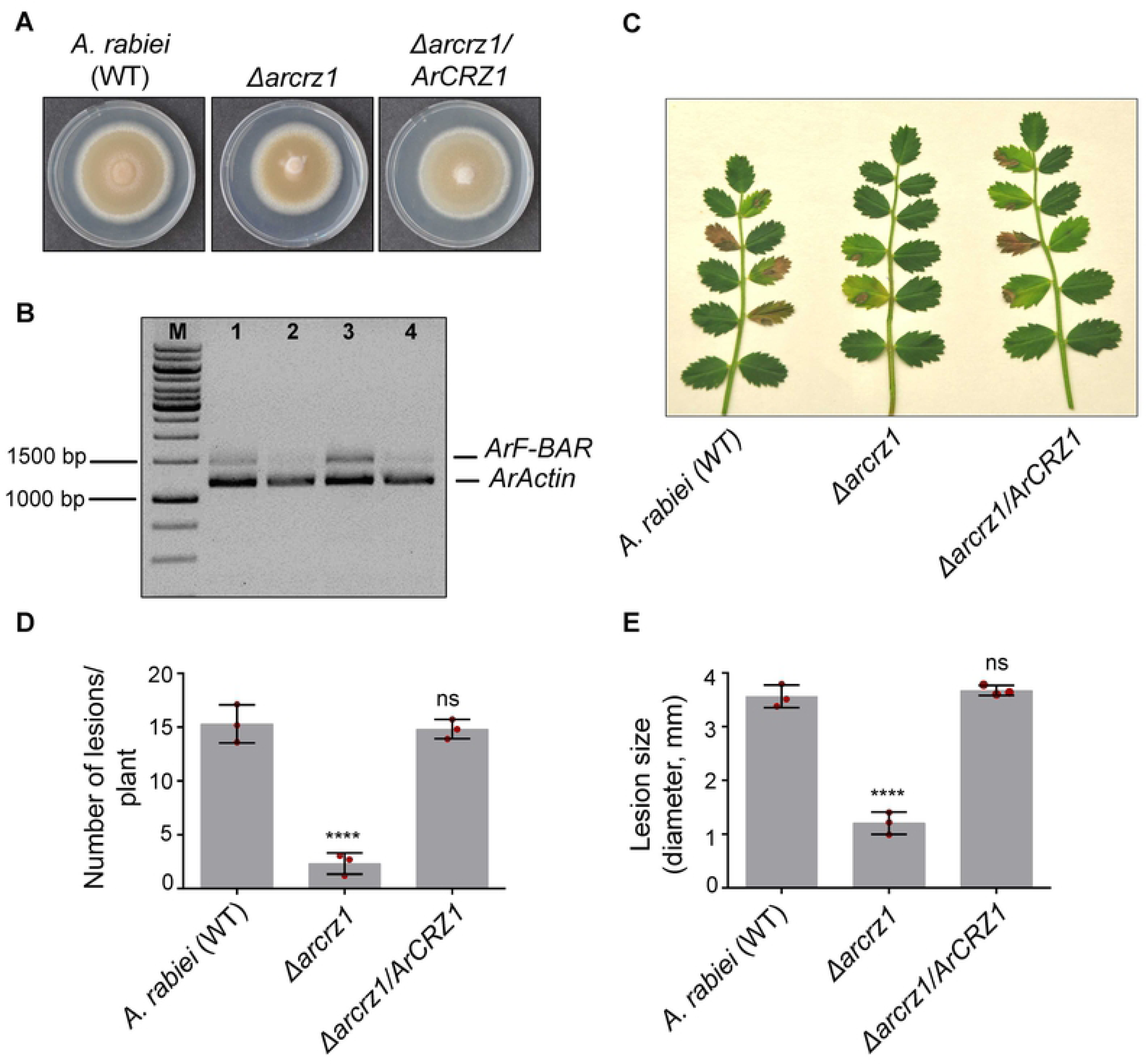
*ArCRZ1* imprint *ArF-BAR* in pathogenicity. (A) Radial growth phenotype of 7 days old *A*. *rabiei* (WT), *Δarcrz1* and *Δarcrz1/ArCRZ1* strains grown on PDA plate. (B) Expression of *ArF-BAR* gene in *Δarcrz1* through semi-quantitative PCR. Lane 1 and 2 showing the *ArF-BAR* expression in *A*. *rabiei* (WT) and *Δarcrz1* while lane 3 and 4 shows the *ArF-BAR* expression in *A*. *rabiei* (WT) and *Δarcrz1*, respectively, in 250 µM menadione treated sample for 0.5 h. (C) Representative image of disease symptoms obtained on AB susceptible chickpea 7 days post conidial inoculation (dpi) of *A. rabiei* (WT), *Δarcrz1* and *Δarcrz1/ArCRZ1* strains. (D) The bar graph showing the number of lesions per plant (WT = 15.31 ± 1.44, *Δarcrz1* = 2.32 ± 0.80 and *Δarcrz1/ArCRZ1* = 14.82 ± 0.73). (E) The bar graph representing the size of the lesions, in diameter (WT = 4.04 mm ± 0.41, *Δarcrz1* = 2.03 mm ± 0.33 and *Δarcrz1/ArCRZ1* = 3.98 mm ± 0.56). The mean and standard deviation (±) were calculated from three biological replicates, counting at least 10 plants for each replicate. These results were quantified using one-way ANOVA, compared with the control (\*\*\*\**p* < 0.0001). Red dots represents the value of each biological replicate/ sample used for the quantitative analysis.

### Loss-of-function of ArCRZ1 compromises pathogenicity similar to *Δarf-bar*

As *ArCRZ1* transcriptionally regulates the expression of *ArF-BAR*, it was hypothesized that the pathogenicity phenotypes of *Δarcrz1* should be similar to those of the *Δarf-bar* mutants. Consistent with this hypothesis, it was observed that *Δarcrz1* mutants displayed compromised pathogenicity during an *in planta* infection bioassay. The number and size of legions were lower in the *Δarcrz1* mutant than in the WT. This pathogenicity defect was rescued in *Δarcrz1/ArCRZ1* (Fig 7C – 7E). The radial growth patterns of *Δarcrz1* mutants grown on PDA supplemented with Ca^2+^, menadione and sodium dodecyl sulfate (SDS) were also monitored. Under these tested stress conditions, the growth inhibition was significantly greater for *Δarcrz1*than for WT. However, the growth phenotype was restored in the *Δarcrz1/ArCRZ1* strain, which suggests that*ArCRZ1* plays a crucial role in calcium ion signaling, oxidative stress response and maintaining cell-wall integrity during infection (S13 and S14 Fig). Taken together, these results demonstrate that ArCRZ1 is a key regulator of the endocytic process. ArCRZ1 induces the expression of *ArF-BAR* by directly binding to its gene promoter region and is critical for *ArF-BAR*-dependent pathogenesis.

## Discussion

Dynamic membrane remodeling is essential for maintaining the integrity and identity of cells and cellular compartments [40]. Biological macromolecules, such as BAR superfamily proteins, can sense or induce membrane curvature. They are well-known modulators of transient membrane deformation [39]. Emerging evidence strongly suggests that EEs are crucial for long-distance intracellular communication. Thus, EEs and their roles have broad implications for a wealth of cellular processes such as growth, development, and virulence in filamentous fungi [4,41,15]. The results of the present study further corroborate this conclusion and provide new information regarding signaling and transcriptional control of F-BAR proteins in phytopathogenic fungi. To date, our understanding of the role played by F-BAR proteins in membrane curvature generation and efficient long-range endosome trafficking in fungi during plant-pathogen interactions remains very limited.

This study unravels the intrinsic mechanism of EE formation in phytopathogenic fungi during polarized hyphal tip growth and host-penetration. To the best of our knowledge, this study provides the first evidence that an F-BAR domain protein can act as a key mandate for pathogenicity. In phytopathogenic fungi, *Ar*F-BAR navigates endosome trafficking via the coordinated action of cell membrane remodeling and actin reorganization. The present findings suggest a model in which *Ar*F-BAR modulates the invagination step and recruits the apical plasma membrane for EE formation during filamentous growth and host penetration of *A. rabiei*. Based on the evidences provided in the present study, we propose that *Ar*F-BAR performs two different but well-coordinated functions that result in EE formation during *A. rabiei*hyphal growth (Fig 8).

**Fig 8.**
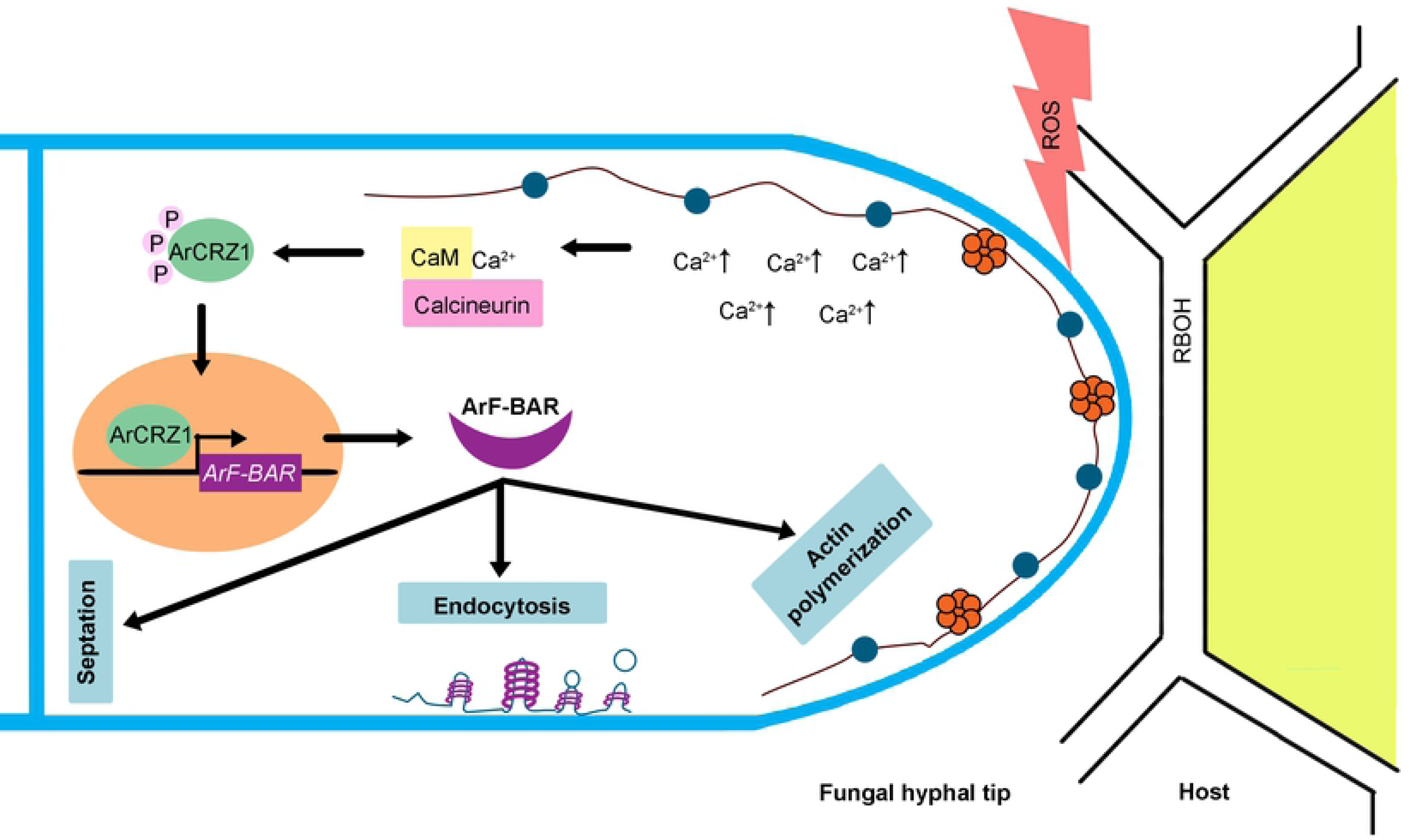
Functional regulation and reprogramming of *ArF-BAR*, required for fungal pathogenicity. On encounter with the pathogen, plant generates ROS at the recognition site. The perception of ROS by the pathogen, results in the upregulation cytosolic Ca^2+^ within the pathogen. Here, in *A*. *rabiei*, increased Calcium level, sensed by calmodulin mediates the activation of calcineurin. Activated calcineurin dephosphorylates ArCRZ1 present in cytoplasm (phosphorylated form of ArCRZ1 remains inactive). Dephosphorylated ArCRZ1 translocates within the nucleus where it regulates the expression of *ArF-BAR*. Further, during polarized growth this ArF-BAR gets localized to hyphal tip where it leads to generation and stabilization of membrane curvature crucial for endosome formation. Additionally, this ArF-BAR protein mediates actin organization and helps in septa formation.

Previous studies, largely conducted in animal models, have highlighted that the F-BAR domain is a membrane-deforming module and is involved in endocytosis [42, 29]. The endocytic event is crucial for the uptake of signal cues and nutrition from the host, and aids apical recycling of membrane receptors and proteins. This process thus helps to maintain the overall polarity of the hyphae that is required for fungal growth and virulence [6, 38]. The generation of EEs and their trafficking involves microtubule dynamics, actin cytoskeleton rearrangements and most importantly, extensive membrane remodeling [43]. Actin dynamics and microtubule organization have been extensively studied. However, the detailed mechanism underlying the functional regulation of fungal EE biogenesis and EE trafficking at the hyphal tip during plant pathogenesis remains poorly understood.

The present study identifies an unprecedented role for the ArF-BAR protein as an essential component of endocytosis that positively regulates EE biogenesis. Unveiling other networks associated with this system, ArF-BAR in coordination with Arp2/3-WASp, was found to mediate actin cytoskeleton assembly at the hyphal tip. The Arp2/3-WASp assembly is a prerequisite for host penetration [44, 45]. The distribution of the F-actin network, which serves as the molecular track for endosome trafficking [8, 35], was disorganized in the *Δarf-bar* mutant, affecting endocytic transport. This phenomenon would explain why the loss of the ArF-BAR function leads to the attenuation of virulence compared to WT. This loss in virulence is similar to that observed in the rice blast fungus *M. oryzae* and *U*. *maydis* where endocytosis is crucial for the recognition of host partners during the early stages of pathogenic development [7]. Therefore, we conclude that ArF-BAR proteins act as membrane-bound tethers to properly recruit the apical plasma membrane for EE formation during endosome trafficking in *A. rabiei*.

During host-pathogen interactions, the key to successful pathogenesis is to overcome the rigid defense responses of the host. Collectively, the first challenge the pathogen encounters is an oxidative burst at the site of infection, which initiates various signaling cascades in the pathogen that aid its survival. Calcium, an essential secondary messenger, mediates one such signaling cascade [46]. In response to stimuli, the cytosolic Ca^2+^ concentration increases [47] and modulates various Ca^2+^-binding proteins such as calmodulin. The Ca^2+^ and calmodulin complex activates calcineurin. In many eukaryotes, calcineurin is known to regulate the activity of CRZ1, which is usually localized in the cytosol in phosphorylated form. Upon activation, CRZ1 gets relocated to the nucleus. In pathogenic fungi, many of the CRZ1-dependent targets, such as those involved in the maintenance of cell wall integrity, thermo-tolerance, cation homeostasis, azole tolerance and hyphal growth have been identified [39]. However, in the present study *ArF-BAR* was identified as a novel ArCRZ1target. The identification of *ArF-BAR* as a target improves our understanding of the novel regulatory mechanism of endocytosis in filamentous fungi, where *ArF-BAR* functions downstream of ArCRZ1.

The development of septa is an important event during hyphal differentiation that is required for the formation of sexual structures and asexual spores [48]. Septation is comparable to cytokinesis that additionally includes cell separation. A cascade of events is involved in septum biogenesis, which includes assembly of the contractile actomyosin ring (CAR), plasma membrane ingression and cell wall constriction [49]. In fission yeast, Cdc15 and Imp2, and in budding yeast, Hof1 are the major F-BAR domain-containing proteins implicated in the formation of the CAR, and the primary and secondary septa during cytokinesis [50, 51]. Additionally, Cdc15 of U. maydis also implicated for similar phenotype [16]. Therefore, the localization of ArF-BAR along the septum ring indicates that endocytosis is one of the pathways responsible for regulating the development of the septa. Septation is initiated at the definitive size of the hyphae [52]. With the initial recognition and establishment of disease in the host, the pathogen needs to proliferate at an enormously increased rate. Therefore, maintaining proper hyphal architecture and polarity is fundamental for pathogenesis that demands rapid coordinated internalization and recycling events [53]. The fungal mutant *Δarf-bar*, deficient in ArF-BAR protein, displayed a delay in septum formation and displayed decreased virulence in the absence of proper hyphal structure.

*In silico* analysis of F-BAR proteins from various filamentous fungi, revealed the evolution of the conserved protein kinase C1 domain. The PKC proteins are phospholipid- and DAG-dependent kinases involved in various intracellular signaling cascades. Fungal PKC has a serine/threonine kinase domain. In *Saccharomy cescerevisae* and *S. pombe*, the PKC is known to activate MAP kinase signaling that helps in cell wall damage repair and in *C. albicans*, PKC helps to provide osmotic tolerance [54]. Preliminary studies from other filamentous pathogens, such as *Aspergillus fumigates* and *Neurosspora crassa*, suggest the involvement of PKC proteins in cellular integrity maintenance. PKC family members have three regulatory domains C1, C2 and HR1, where the C1 domain of classic PKC enzymes has the ability to bind to DAG and phorbol esters [55]. Studies have reported the presence of two different cysteine-rich (C1) motifs, C1A and C1B, termed as “typical” and “atypical,” respectively, depending on whether they “do” or “do not” bind to DAG and phorbol esters [57]. Interestingly, the presence of a specific C1 domain provides uniqueness to the F-BAR protein of filamentous fungi. However, in the current study, the targeted mutation of this domain did not have any functional relevance for virulence. This lack of relevance may have been for two reasons; a) C1 might be present in an “atypical” form or b) the lack of the C2 domain in ArF-BAR; both the C1 and C2 domains are required for full-enzymatic activation of PKC [58].

In summary, we propose that the ArF-BAR protein of *A. rabiei* has the potential to interactively affect the hyphal growth and pathogenic developmental trajectories of filamentous fungi. This evolving model provides mechanistic insight into the role of a membrane scaffolding protein in the process of endosome trafficking in fungal pathogenesis. In turn, this provides many additional potential targets for the development of effective and durable strategies to control AB fungal disease. Therefore, the observations of the present study in context to the intracellular trafficking, during the early stages of plant-pathogen interactions, have broad relevance for shaping disease-control strategies. These findings may also be helpful for the disease control of animal-infecting fungal pathogens such as *A. fumigatus* and *C. albicans*. Thus, further studies will be directed to characterize the interacting partners influenced by ArF-BAR during endosome formation. A diverse spectrum of studies has revealed the existence of two parallel independent endocytic mechanisms: a) clathrin-mediated and b) clathrin-independent [59]. Similarly, extensive studies will be required to fully elucidate the ArF-BAR-mediated endocytic mechanism. Such studies would help us to understand the complex interplay underlying the endosome formation required for fungal virulence. Further, the understanding of its regulatory mechanism would help in scrutinizing the molecular and cellular basis of disease development, which would subsequently help develop disease-control strategies for filamentous fungi.

## Methods

### Fungal strains and growth conditions

A virulent isolate of *Ascochyta rabiei* (ArD2; ITCC No. 4638) was procured from IARI, New Delhi. The single spore culture of this mating type 2 isolate was generated and maintained as wild type fungi for research work. The WT and its derivative fungal strains (S2 Table) were maintained on potato dextrose agar (PDA; Difco Laboratories, pH 5.2-5.5) at 22°C for 10-15 days to assess the growth pattern and colony characteristics [26]. The fungus was routinely subcultured on the PDA plate supplemented with chickpea extract to maintain virulence. To determine the vegetative growth pattern of fungal mycelia in response to oxidative stress, PDA plate supplemented with menadione (250 µM and 500 µM; Sigma-Aldrich, USA) and H_2_O_2_ (2 mM; Sigma-Aldrich, USA) were used. The conidial suspensions of 10 µl (1x10^3^ conidia/ml) were inoculated at the center of PDA plate for growth assay. After 10 days of incubation under optimum condition, the diameters of fungal colonies were measured using ImageJ software. Three independent biological experiments were performed with three technical replicate each time.

### RNA extraction and expression analysis

Total RNA was extracted from 5-6 days old fungal mycelia grown in potato dextrose broth (PDB; Difco Laboratories, USA) or from plant tissues inoculated with WT using TRIzol reagent (Invitrogen, USA). The isolated total RNA was subjected to DNase1 (Promega, USA) treatment and subsequently used for first-strand cDNA synthesis using SuperScript IV reverse transcriptase (ThermoFisher Scientific). Targeted gene expression was determined by qRT-PCR with ABI7900 (Applied Biosystems, USA), using SYBR Green PCR master mix (Applied Biosystems, USA). Relative expression of fungal genes was calculated after normalized with elongation factor α (*ArEFα*; ST47_g4052) using 2^-ΔΔct^ method [60]. The data were analysed from three biological replicates each having three technical replicates.

### Site-directed mutagenesis

The mutations at requisite sites of *ArF*-*BAR* were achieved by PCR amplification of pET28a(+):*ArF-BAR* clone with pre-designed primers containing mutations of interest. Mutagenesis was performed using the QuikChange II Site-directed mutagenesis kit (Agilent, USA). Web-based Quik Change Primer Design tool (www.agilent.com/genomics/qcpd) was used to design primers. The presence of mutations in clones was confirmed by Sanger sequencing.

### Targeted gene knockout and complementation in *A*. *rabiei*

Homologous gene replacement with *hph* cassette strategy was used to generate knockout (KO) constructs for *A. rabiei* genes. Genomic DNA was isolated from 5-day grown PDB culture of *A. rabiei* using GenElute^TM^ Plant Genomic DNA miniprep kit (Sigma-Aldrich, USA). The 5′ flanking genomic sequences of *ArF-BAR* were amplified from *A. rabiei* genomic DNA using primer pairs of ArF-BARKOif5F and ArF-BARKOif5R while the 3′ flanking sequences were amplified using Ar72KO3 and Ar72KO4 (S3 Table). These amplified 5′ and 3′ flanking sequences were cloned sequentially into pGKO2 vector at *Kpn*I/*Pst*I and *Bam*HI/*Eco*RI sites, respectively. The cloned *ArF*-*BAR* gene replacement cassette of ∼3.4 kb, including 5′ and 3′ flanking sequence along with *hph*, was amplified using primer pair ArF-BARKOif5F and Ar72KO4, and transformed into *A. rabiei* protoplasts. The *A. rabiei* protoplast transfection was performed as described earlier [61]., with some minor modifications. The putative transformants were selected on a PDA plate supplemented with 50 µg/µl hygromycin. To generate the complementation constructs of *ArF-BAR* and its mutated (mut1 and mut2) versions, about 4.4 kb DNA fragment having the native promoter, ORF region, and TrpC terminator was amplified and cloned into pBIF2 vector (Bacterial selection-kanamycin; Fungal selection-G418) at *Eco*RI and *Hind*III sites. These three constructs of *ArF*-*BAR* were independently transformed into *Δarf-bar* mutant strain by ATMT [27]. Similarly, *A*. *rabiei ArCRZ1* gene knockout mutant and complementation strains were developed.

### Gene knockout confirmation by PCR and Southern blot

The single spore culture of putative KOs selected on hygromycin was initially screened by genomic PCR. A primer set binding at position 5′ to the homologous recombination region and TrpC promoter was used for gene upstream region while a primer set binding at *Hph* gene and position 3′ to the homologous recombination region was used for gene downstream integrity check (S3 Table). The complemented strains single spore culture was also confirmed by genomic PCR. The PCR positive *Δarf-bar*, and *Δarcrz1* mutants (KOs) and complemented strains were further verified by Southern blot. The genomic DNA of WT, *Δarf-bar* and *Δarcrz1* mutants was digested with *Eco*RI enzyme while genomic DNA was digested with *Eco*RI and *Hind*III for the complemented mutant strains (*Δarf-bar/ArF-BAR*, *Δarf-bar/ArF-BAR^mut1^*, *Δarf-bar/ArF-BAR^mut2^*, and *Δarcrz1/ArCRZ1*). The digested DNA was separated along with λ DNA/*Hind*III marker (ThermoFisher Scientific) and blotted to a membrane followed by hybridization with radioactive probe [62] prepared using the random primers labelingNEBlot^®^ kit (New England Biolabs, USA). The band detection was carried out using Typhoon^®^ phosphor imager (GE Healthcare, USA).

### Pathogenicity assay

Two-week-old susceptible chickpea (Pusa 362) plants grown in plant growth chambers under controlled conditions (D/N temperature: 24°C/18°C; Relative Humidity: 80%; light intensity 250 µE/m^2^/s; D/N light duration: 14/10 h) were used for infection assays. Conidial suspensions of *A*. *rabiei* strains were collected separately from a 20-days-old PDA plate grown culture. Two-weeks-old plants were spray inoculated with conidial suspensions diluted to 2x10^6^ conidia/ml.

Plants were again kept under optimum conditions. Disease lesions were examined 5-7 days after spray inoculation.

### *In vitro* Protein purification

Respective cDNAs were cloned in pET28a (+) and transformed into *E. coli* BL21-CodonPlus (DE3)-RIPL cells. The protein expression was induced with 0.5 mM Isopropyl β-D-thiogalactoside (IPTG) for 6 h at 23°C. Bacterial pellet was lysed in buffer [500 mMNaCl, 50 mM NaPO_4_ (pH 8.0), 10 mM Imidazole, 1mM β-Mercaptoethanol, 1 mg/ml lysozyme and 1mM Phenylmethylsulfonyl fluoride (PMSF)] by incubating 30 min on ice, followed by sonication. The cell lysates were precipitated followed by 0.45 µm filtration. The cleared lysate was incubated with Ni-NTA resin for 30 min at 4°C. The His-tagged fusion protein was purified using a Ni-NTA column (Bio-Rad, USA). Protein was eluted in elution buffer [500 mMNaCl, 50 mM NaPO_4_ (pH 8.0), 10% glycerol and 200 mM imidazole]. The quality and quantity of eluted protein were checked by SDS/PAGE and Bradford assay, respectively. The proteins were dialyzed in respective compatible buffers followed by concentration.

### Liposome preparation and Tubulation assay

Liposomes were prepared as described previously [63] using 70% Phosphatidylethanolamine (POPE), 20% Phosphatidylcholine (POPC), and 10% Rhodamine B-conjugated PE (Echelon Biosciences, USA). In an amber vial, all the lipids were initially dissolved in chloroform: methanol (65:35;v/v) mixture and vial were kept under liquid nitrogen for 10 min before being immediately subjected to vacuum desiccation/lyophilisation for 2 h at 60 mTorr. The lipids were hydrated with buffer [25mM Tris-HCl (pH 6.8) and 100mM NaCl] and subjected to three freeze-thaw cycles of 5 min each at 68 °C and liquid nitrogen. Extrusion was performed at 68 °C on a pre-heated mini extruder (Avanti Polar Lipids, USA). The prepared liposomes were diluted as desired and immediately proceeded for tubulation assay. Before use, the purified proteins were subjected to 100,000 g centrifugation for 20 min at 4 °C to remove the aggregate proteins. To examine tubule formation, the mixed liposome and protein samples were analysed in Lumox^®^24-well plate (Millipore, USA) by live-cell imaging on Axio Examiner.Z1 (Zeiss microscope).

### Actin polymerization assay

The actin polymerization modulation activity of the proteins was checked using Actin Polymerization Biochem Kit (Cytoskeleton, USA), using the manufacturer’s instruction. Freshly solubilized components; 13 nM Arp2/3 protein complex and 15 nMWASp-VCA domain-GST purified (Cytoskeleton, USA) along with freshly purified ArF-BAR protein in concentrations of 200, 400 and 600 nM were used. The reaction was carried in an opaque 96-well plate and kept in the dark. The actin polymerization rate was recorded by monitoring the pyrene fluorescence signals using CLARIO^®^ star plate reader (BMG Labtech, Germany) with the following settings; slow kinetics, 60 sec interval, λ_ex_ = 360±15 nm and λ_em_ = 420±20 nm.

### Liposome co-sedimentation assay

The purified fusion proteins were pre-centrifuged, before the assay, at 100,000 g for 15 min to remove protein aggregates. Protein from the supernatant was mixed with freshly prepared synthetic liposomes with gentle tapping. Ultracentrifugation was performed at 100,000 g for 15 min at 4°C and the supernatant and the pellet were carefully separated. The supernatant was mixed with 1:1 loading buffer while the pellet was re-suspended in 2x loading buffer. Samples were analysed on Coomassie stained SDS-PAGE.

### Yeast two-hybrid assays

The interactions of various combinations between ArF-BAR, ArActin, ArWASP, and ArF-BAR domains in yeast cytoplasm were examined using the split-ubiquitin based DUALhunter system (Dualsystems Biotech). The ORFs were cloned at *Not*I and *Asc*I restriction sites or by LR clonase II into pGDHB1 and pGPR3-N vectors. The cloned plasmids were co-transformed along with necessary controls into NMY51 strain using the EZ-Yeast transformation kit (MP Biomedicals, USA) and plated on SD-L/-W plates. The plating of yeast clones on required synthetic media to check protein-protein interactions in yeast was done as described previously [22]. All the interactions were verified by three independent experiments.

### Electrophoretic mobility shift assay (EMSA)

His-tagged ArCRZ1 (*His*-*ArCRZ1*) expression construct was developed and recombinant protein was purified. The wild type and mutated CDRE DNA fragments were assembled by annealing oligonucleotide pairs in a thermal cycler by heating at 95° C for 5 min followed by cooling at RT for 15 min. End labelling of DNA fragments was performed by [γ^32^P] dATP BRIT, India), and Polynucleotide Kinase (Thermo Fisher Scientific, USA). Additionally, for competition assays, two complementary oligonucleotides were annealed at equimolar concentration. Purified His-ArCRZ1 protein was incubated with 10 ng of labelled DNA fragment in the presence of 1 mg of poly-deoxy-inosinic-deoxy-cytidylic acid [poly (dI-dC)] and 1X binding buffer (15mM HEPES (pH 7.6), 0.2 mM MgCl_2_, 35 mMKCl, 1 mM DTT and 1% glycerol) in a reaction volume of 30 µl for 25 min at room temperature. DNA loading dye was used to terminate the reaction. The competitive assays were performed using 50, 100 and 200 times of specific fragments in excess. To identify the relative binding, the complexes were resolved on 6% native PAGE, dried, and autoradiographed on X-ray films.

## Microscopic methods

### Sample preparation for microscopic analysis

The conidia from the respective fungal strains were harvested in 1 ml sterile distilled water from 15-day-old fungal mycelia grown on PDA plates. The conidial suspension was filtered through Mira cloth. Ten microlitres (1×10^6^ conidia/ml) of suspension was kept on a sterile glass coverslip and allowed to grow under the optimal condition for 24 h in dark. To investigate the localization of chimeric proteins in fungal hyphae, growing on chickpea stem peel, the conidia were allowed to grow for 36 h under optimum conditions of infection. The grown hyphae samples were then used for confocal laser scanning microscopy.

### Confocal laser scanning microscopy

For microscopic studies, TCS SP5 and TCS SP8 confocal laser scanning microscope (Leica Microsystems, Germany) were used. For subcellular localization, the conidia were harvested from transgenic fungal strains expressing fluorescent-tagged proteins. The Z-stacked images with 1 µm step size were acquired using a high-resolution CCD camera. For calcofluor-white (CW), the hyphae grown on the glass slide were incubated for 10 min in the CW solution (Sigma-Aldrich, USA). After incubation, the stained hyphae were rinsed with PBS (pH 7.4) followed by rinsing with sterile distilled water, before image acquisition. For FM4-64 uptake, 1 ml of harvested conidial suspension was centrifuged at 2,500 g, washed twice with sterile distilled water and then allowed to germinate on glass slides. The aqueous solution (10 µM) of FM4-64 dye (Invitrogen, USA) was added directly to the fungal mycelia. After 10 min incubation, FM4-64 dye was rinsed from slides thoroughly before imaging fungal hyphae. Images were captured in TCS SP8.

For host penetration assay, infected chickpea leaves (WT and *Δarf-bar*) were placed in 100 % ethanol for 48 h to undergo bleaching for the complete removal of chlorophyll.

Subsequently, leaves were incubated for 4 h in 10 % KOH at RT followed by washing 4-5 times in Phosphate buffer saline (PBS, pH 7.4). The processed leaves were then stained with chitin specific dye WGA-AF 488 (Invitrogen, USA). The leaf samples were rinsed in PBS (pH 7.5) before microscopic visualization. Confocal images were captured on a TCS SP5 confocal microscope.

For sub-cellular localization of ArCRZ1::EYFP under oxidative stress condition, chimeric protein-expressing fungal conidia were isolated and allowed to grow for 12 h. The hyphae were exposed to CaCl_2_ (200 mM, Sigma-Aldrich, USA) for 1 min prior to microscopy. To assess the involvement of calcineurin in nuclear localization of ArCRZ1::EYFP, the hyphae was exposed to 5 µg/µl FK506 (Sigma-Aldrich, USA), 5 min prior to the addition of CaCl_2._ The confocal images were acquired in TCS SP5.

For subcellular localization of ArCRZ1::EYFP during infection, susceptible plants were challenged with conidia expressing ArCRZ1::EYFP. The images were acquired after 48 h using TCS SP5.

### Quantification and Statistical analysis

Quantification analysis of relative fluorescent intensity, lesion size, radial diameter, the distance of septa from the hyphal tip and distance travelled were analysed by ImageJ/Fiji software. To calculate the significance of means/differences between two groups, Student’s t-test and oneway

ANOVA followed by Tukey test between multiple groups were performed using GraphPad Prism 6. Significance was accepted at *p* < 0.05, as noted in the text of legends. Replicates are indicated in the legends.

### Bioinformatic analysis

All the gene and protein sequences were acquired from NCBI server (http://www.ncbi.nlm.nih.gov). Multiple sequence alignment was performed by Praline search (http://www.ibi.vu.nl/programs/pralinewww/). Phylogenetic analysis was performed with MEGA7.0.21 software. Distinctive domain organization of the protein was determined by SMART search (http://smart.embl-heidelberg.de/). The putative TFs binding sites were identified by YEASTRACT-DISCOVERER Database (http;//yeastract.com). The theoretical pI and molecular weight of the chimeric proteins were determined by Expasy compute pI/Mw tool (http://web.expasy.org/cgi-bin/compute_pi/pi_tool). Protein IDs of protein sequences used in this study are: ArF-BAR: KZM20872.1, ArRab5: KZM19760.1, ArRab7: KZM26450.1, ArActin: KZM21342.1, ArWASP: KZM19192.1 and ArCRZ1: KZM25117.1.

## Acknowledgements

We gratefully acknowledge a research grant from Department of Biotechnology, Government of India (File No: BT/PR10605/PBD/16/791/2008 and BT/AGR/CG-Phase II/01/2014) and a core grant from National Institute of Plant Genome Research, New Delhi, India for funding this work. MS, AS, KS and KK acknowledge Council of Scientific and Industrial Research and University Grants Commission, Government of India for their fellowships.

## Author contributions

PKV and VK designed the project; MS, AS and KS carried out experiments; PKV, KK and KS supervised the work; all authors analysed and discussed the data; MS, AS, KK and VK wrote the manuscript; all the authors reviewed and approved the manuscript; MS and AS contributed equally to this work.

## SUPPORTING INFORMATION

**S1 Fig**

**Maximum likelihood phylogeny of F-BAR sequences**.

Sequences from *Ascochyta rabiei*, *Parastagonospora nodorum*, *Alternaria alternate*, *Bipolaris zeicola*, *Bipolaris victoriae*, *Bipolaris oryzae*, *Bipolaris sorokiniana*, *Bipolaris maydis*, *Sclerotinia sclerotiorum*, *Botrytis cinerea*, *Marssonina brunnea*, *Histoplasma capsulatum*, *Blastomyces dermatitidis*, *Blastomyces gilchristii*, *Emmonsia crescens*, *Paracoccidioides brasiliensis*, *Uncinocarpus reesii*, *Coccidioides immitis*, *Aspergillus nidulans*, *Aspergillus rambellii*, *Aspergillus ochraceoroseus*, *Aspergillus kawachii*, *Aspergillus lacticoffeatus*, *Aspergillus oryzae*, *Aspergillus flavus*, *Aspergillus parasiticus*, *Aspergillus nomius*, *Aspergillus terreus*, *Aspergillus fumigatus*, *Aspergillus lentulus*, *Aspergillu sclavatus*, *Aspergillus fischeri*, *Trichoderma reesei*, *Cordyceps confragosa*, *Fusarium oxysporum*, *Fusarium verticillioides*, *Fusarium fujikuroi*, *Fusarium graminearum*, *Fusarium pseudograminearum*, *Fusarium langsethiae*, *Neonectria ditissima*, *Metarhizium anisopliae*, *Metarhizium acridum*, *Pyricularia grisea*, *Pyricularia oryzae*, *Verticillium dahlia*, *Neurospora crassa*, *Drosophila melanogaster*, *Schizosaccharomyces pombe*, *Phytophthora sojae*, *Phytophthora graminis*, *Phytophthora striiformis*, *Ustilago maydis*, *Rhizopus delemar*, *Cryptococcus*, *Phytophthora infestans*, Yarrowialipolytica, *Kluyveromyces lactis*, Naumovozymacastellii, *Saccharomyces cerevisiae*, *Candida glabrata*, Eremotheciumgossypii, Clavisporalusitaniae, Meyerozymaguilliermondii, Candida parapsilosis, *Candida tropicalis*, *Candida dubliniensis*, *Candida albicans*, Ganodermalucidum, and *Homo sapiens*.

The multiple sequence alignment of protein was performed by PROMALS3D software and the phylogeny was constructed using a software MEGA7.0.21. The bootstrap values, derived from 1000 iterations, validated the obtained phylogeny.

**S2 Fig**

**Conserved nature of ArF-BAR protein.**

Multiple sequence alignment showing the conservation of ArF-BAR protein with *S. cerevisiae* BZZ1p, *Drosophila* Cdc42-interacting protein 4 (CIP4) and *Drosophila* Syndapin proteins. Colour code for sequence conservation varies from blue (least conserved) to red (highly conserved). The alignment of the protein is determined by Praline software using default parameters. The black box marks the presence of positively charged residues of F-BAR domain. Asterisk (*) represents the residues in PKC domain required for interaction with DAG.

**S3 Fig**

**Expression and purification of ArF-BAR protein in *E. coli*.**

(A) His purification of bacterially expressed ArF-BAR protein. Analysis of the purification of recombinant ArF-BAR as shown by SDS-PAGE. (B) Intense tubular network in synthetic liposomes is formed after 30 min incubation with purified recombinant *Ar*F-BAR protein. Inset showing the enlarged view of dense tubular network originating from a liposome. (C-D) His purification of ArF-BAR^mut1^, ArF-BAR^mut2^. The protein was visualised by Comomassie Brilliant Blue staining. (UI, crude extract of un-induced samples after centrifugation; I, crude extract of induced samples after centrifugation; FT, flow-through fraction of the Nickel chelating resin column; W5- 5^th^ wash fraction of the Nickel chelating resin column; E1, E2 and E3 eluate fractions of the Nickel chelating resin column showing the purified ArF-BAR protein). Protein standards are shown (M), and their masses are indicated in kDa.

**S4 Fig**

**Southern confirmation for the successful replacement of *ArF-BAR* gene with Hygromycin cassette and its complementations.**

(A) The schematic representation of *A*. *rabiei* knockout mutant generation by the homologous recombination approach to obtain targeted *ArF-BAR* gene deletion mutants (*Δarf-bar*). The bar represents the genomic region used to generate probe for Southern confirmation. (B) The representative Southern blot confirming successful *ArF-BAR* gene deletion (*Δarf-bar*), with single integration of *hph* at replacement site. Along with the confirmation of *ArF-BAR* complementation in *Δarf-bar,* followed successful generation of *Δarf-bar/ArF-BAR*^mut1^and *Δarf-bar/ArF-BAR*^mut2^ complementations.

**S5 Fig**

**The schematic representation of constructs.**

(A, B, C) Constructs used to generate different *Δarf-bar* mutant complemented strains (*Δarf-bar/ArF-BAR, Δarf-bar/ArF-BAR*^mut1^ and *Δarf-bar/ArF-BAR*^mut2^) under the control of native promoter of *ArF-BAR* gene.

**S6 Fig**

**Disease symptoms on AB susceptible chickpea plants 10 dpi.**

The susceptible plants inoculated with conidia of *A. rabiei* (WT), *Δarf-bar/ArF-BAR* and *Δarf-bar/ArF-BAR*^mut2^, showed severe disease symptoms with increasing duration of infection. *Δarf-bar and Δarf-bar/ArF-BAR*^mut1^ challenged plants were healthier even after 10 dpi.

**S7 Fig**

**Evaluation of chickpea host cells penetration by *A. rabiei* (WT) and *Δarf-bar* strains growing hyphae.**

(A, B) Confocal images showing the depth of penetration 48 hpi by *A. rabiei* (WT) and *Δarf-bar* strains, respectively, in AB susceptible chickpea leaves. Fungal hyphae were stained with WGA-488 for visualization, prior to microscopy. The *Z*-stacked images were acquired till 23 µm depth, starting from the surface of the leaves. The image is the representation of maximum projections of all the Z-stacks. Scale bar = 5 µm.(C) The bar graph, representing mean and SD, shows the difference in ability to penetrate within the host by *A. rabiei* (WT) and *Δarf-bar.* The results were quantified using Student’s t-test one tailed compared to its control (\**p* < 0.0079).

**S8 Fig**

**Colony morphology of *Δarf-bar* and mutant complemented strains, under oxidative stress conditions.**

Colony morphology and growth assay of *A. rabiei* (WT), *Δarf-bar* mutant and mutant complemented strains, observed after 10 days after incubation at 22°C. (A) PDA is supplemented with 250 µM and 500 µM menadione, and 2mM H_2_O_2_ to induce oxidative stress. *Δarf-bar* and *Δarf-bar/ArF-BAR*^mut1^ exhibited extreme sensitivity towards oxidative stress condition as compared to *A. rabiei* (WT). (B) The graph represents mean and SD values of fungal strains radial diameter in the presence of oxidative stress. All the growth assays were performed in triplicate. The results were quantified using two-way ANOVA, Tukey’s multiple comparisons. \*\*\*\**p* < 0.0001, \*\*\**p* < 0.001, \*\**p* < 0.01, \**p* < 0.05, ns = non-significant. Red dots represents the value of each biological replicate/ sample used for the quantitative analysis.

**S9 Fig**

**Distribution of ArF-BAR::EYFP during *in planta* infection.**

Confocal micrographs showing the punctate distribution of ArF-BAR::EYFP during host infection. The representative image is the maximum intensity projection of all Z-stack images with 0.5 µm step size, acquired after 48 hpi of susceptible chickpea with fungal conidia expressing chimeric ArF-BAR::EYFP. Scale bar = 5 µm (n= 12).

**S10 Fig**

**Conserved nature of F-BAR to form homodimer.**

(A) Interaction analysis of *Ar*F-BAR proteins by yeast two-hybrid (Y2H). Split-Ubiquitin based Y2H system was used to determine the homodimerization between ArF-BAR protein. Plates were photographed after 48 h of yeast growth. Strong positive interaction between two ArF-BAR proteins was reflected with the growth on QDO (SD/-L/-W/-A/-H) media and X-gal overlay assay to check the activation *LacZ* gene. (B) The *Ar*F-BAR does not localizes with the late endosome as ArF-BAR::mCherry and ArRab7::EYFP failed to co-localize. The representative image is the maximum intensity projection of all z-stack images with 1 µm step size. Images were acquired after 12 h post-germination of fungal conidia expressing ArF-BAR::mCherry and ArRab7::EYFP. Scale bar = 5 µm, (n= 5).

**S11 Fig**

**F-BAR domain of ArF-BAR is not the direct member to interact.**

(A) The yeast two-hybrid result showing the positive interaction of ArF-BAR protein with ArWASp. (B) F-BAR domain (1-325 amino acids) of ArF-BAR protein [F-BAR(_ArF-BAR_)] failed to interact with ArWASp in Y2H system. Plates were photographed 48 h after yeast spotting. The interaction was confirmed through three independent replicates.

**S12 Fig**

**Nuclear localization of ArCRZ1 during *in planta* infection.**

(A) Schematic representation of domain organisation of ArCRZ1 protein. (B) Confocal images showing the nuclear localization of ArCRZ1, during *in planta* fungal growth. Confocal images were acquired 48 hpi of AB susceptible chickpea with conidia of strain expressing ArCRZ1::EYFP. Scale bar = 5 µm, (n= 30). (C) The schematic map showing the homologous recombination based knockout approach used for targeted Ar*CRZ1* gene deletion mutant (*Δcrz1*) strain generation. (D) Schematic representation of *Δarcrz1/ArCRZ1* complementation construct under the native promoter of *ArCRZ1*. The genomic region used to generate probe for Southern confirmation is being highlighted. (E) The Southern blot result confirmed successful *ArCRZ1* gene deletion (*Δarcrz1*), with single integration of *hph* at replacement site. Along with the complementation confirmation of *ArCRZ1* in *Δarcrz1* mutant.

**S13 Fig**

**Growth phenotypes of *A. rabiei* (WT), *Δarcrz1* and *Δarcrz1/ArCRZ1* under various stress conditions**

Colony morphology and growth assay of *A. rabiei* (WT), *Δarcrz1* mutant and *Δarcrz1/ArCRZ1* complementation strains observed under various stress conditions after 10 days incubation at 22°C. PDA is supplemented with 250 µM menadione and 2mM H_2_O_2_, 0.07 M, 0.2 M, 0.4 M CaCl_2_ and 0.01% SDS. The *Δarcrz1* mutant exhibited extreme sensitivity towards various stress conditions as compared to *A. rabiei* (WT). The mutant strain was highly sensitive towards CaCl_2_ and fails to grow at higher concentration of 0.4 M.

**S14 Fig**

**Radial diameter *A. rabiei* (WT) and *Δarcrz1* mutant and *Δarcrz1/ArCRZ1* complementation strains**

The bar graph represents mean and SD of radial diameter for the fungal strains in the presence of various stresses. All the growth assays were performed in triplicates. The results were quantified using two-way ANOVA, Tukey’s multiple comparisons. The statically significant differences are shown with \*\*\*\**p* < 0.0001, \*\*\**p* < 0.001, \*\**p* < 0.01, \**p* < 0.05, ns = non-significant. Red dots represents the value of each biological replicate/ sample used for the quantitative analysis.

**S1 Table**. YEASTRACT result for the putative transcription factors bindings on the upstream regulatory sequences of ArF-BAR.

**S2 Table**. *Ascochyta rabiei* strains used in this study.

**S3 Table**. List of oligonucleotides used in this study.

## References

1. Takeshita N, Evangelinos M, Zhou L, Serizawa T, Somera-Fajardo RA, Lu L, et al. Pulses of Ca^2+^ coordinate actin assembly and exocytosis for stepwise cell extension. Proc Natl Acad Sci U S A. 2017;114: 5701–5706. doi:10.1073/pnas.1700204114

2. Riquelme M, Aguirre J, Bartnicki-García S, Braus GH, Feldbrügge M, Fleig U, et al. Fungal Morphogenesis, from the Polarized Growth of Hyphae to Complex Reproduction and Infection Structures. Microbiol Mol Biol Rev. 2018. doi:10.1128/mmbr.00068-17

3. Etxebeste O, Espeso EA. Neurons show the path: Tip-to-nucleus communication in filamentous fungal development and pathogenesisa. FEMS Microbiology Reviews. 2016. doi:10.1093/femsre/fuw021

4. Bielska E, Higuchi Y, Schuster M, Steinberg N, Kilaru S, Talbot NJ, et al. Long-distance endosome trafficking drives fungal effector production during plant infection. Nat Commun. 2014. doi:10.1038/ncomms6097

5. Cell Biology of Hyphal Growth. The Fungal Kingdom. 2017. doi:10.1128/microbiolspec.funk-0034-2016

6. Higuchi Y, Shoji JY, Arioka M, Kitamoto K. Endocytosis is crucial for cell polarity and apical membrane recycling in the filamentous fungus *Aspergillus oryzae*. Eukaryot Cell. 2009;8: 37–46. doi:10.1128/EC.00207-08

7. Takeshita N. Coordinated process of polarized growth in filamentous fungi. Biosci Biotechnol Biochem. 2016;80: 1693–1699. doi:10.1080/09168451.2016.1179092

8. Takeshita N, Higashitsuji Y, Konzack S, Fischer R. Apical sterol-rich membranes are essential for localizing cell end markers that determine growth directionality in the filamentous fungus *Aspergillus nidulans*. Mol Biol Cell. 2008. doi:10.1091/mbc.E07-06-0523

9. Rao Y, Rückert C, Saenger W, Haucke V. The early steps of endocytosis: From cargo selection to membrane deformation. European Journal of Cell Biology. 2012. doi:10.1016/j.ejcb.2011.02.004

10. Frost A, Perera R, Roux A, Spasov K, Destaing O, Egelman EH, et al. Structural Basis of Membrane Invagination by F-BAR Domains. Cell. 2008. doi:10.1016/j.cell.2007.12.041

11. Habermann B. The BAR-domain family of proteins: a case of bending and binding? The membrane bending and GTPase-binding functions of proteins from the BAR-domain family. EMBO Rep. 2004. doi:10.1038/sj.embor.7400105

12. Krauss M, Haucke V. A novel twist in membrane dephormation. Developmental Cell. 2014. doi:10.1016/j.devcel.2014.09.016

13. Saarikangas J, Zhao H, Pykäläinen A, Laurinmäki P, Mattila PK, Kinnunen PKJ, et al. Molecular Mechanisms of Membrane Deformation by I-BAR Domain Proteins. Curr Biol. 2009. doi:10.1016/j.cub.2008.12.029

14. Fuchs U, Hause G, Schuchardt I, Steinberg G. Endocytosis is essential for pathogenic development in the corn smut fungus *Ustilago maydis*. Plant Cell. 2006;18: 2066–2081. doi:10.1105/tpc.105.039388

15. Zimmerberg J, Kozlov MM. How proteins produce cellular membrane curvature. Nature Reviews Molecular Cell Biology. 2006. doi:10.1038/nrm1784

16. Böhmer C, Ripp C, Bölker M. The germinal centre kinase Don3 triggers the dynamic rearrangement of higher-order septin structures during cytokinesis in *Ustilago maydis*. Mol Microbiol. 2009. doi:10.1111/j.1365-2958.2009.06948.x

17. Lin L, Chen X, Shabbir A, Chen S, Chen X, Wang Z, et al. A putative N-BAR-domain protein is crucially required for the development of hyphae tip appressorium-like structure and its plant infection in *Magnaporthe oryzae*. Phytopathol Res. 2019;1: 1–15. doi:10.1186/s42483-019-0038-2

18. Dagdas YF, Yoshino K, Dagdas G, Ryder LS, Bielska E, Steinberg G, et al. Septin-mediated plant cell invasion by the rice blast fungus, *Magnaporthe oryzae*. Science (80-). 2012;336: 1590–1595. doi:10.1126/science.1222934

19. Soulard A, Lechler T, Spiridonov V, Shevchenko A, Shevchenko A, Li R, et al. *Saccharomyces cerevisiae* Bzz1p Is Implicated with Type I Myosins in Actin Patch Polarization and Is Able To Recruit Actin-Polymerizing Machinery In Vitro. Mol Cell Biol. 2002. doi:10.1128/mcb.22.22.7889-7906.2002

20. Fricke R, Gohl C, Dharmalingam E, Grevelhörster A, Zahedi B, Harden N, et al. Drosophila Cip4/Toca-1 Integrates Membrane Trafficking and Actin Dynamics through WASP and SCAR/WAVE. Curr Biol. 2009. doi:10.1016/j.cub.2009.07.058

21. Arasada R, Pollard TD. Distinct roles for F-BAR proteins Cdc15p and Bzz1p in actin polymerization at sites of endocytosis in fission yeast. Curr Biol. 2011. doi:10.1016/j.cub.2011.07.046

22. Kumar K, Purayannur S, Kaladhar VC, Parida SK, Verma PK. mQTL-seq and classical mapping implicates the role of an AT-HOOK MOTIF CONTAINING NUCLEAR LOCALIZED (AHL) family gene in Ascochyta blight resistance of chickpea. Plant Cell Environ. 2018;41: 2128–2140. doi:10.1111/pce.13177

23. Pandey BK, Singh US, Chaube HS. Mode of Infection of Ascochyta Blight of Chickpea Caused by *Ascochyta rabiei*. J Phytopathol. 1987. doi:10.1111/j.1439-0434.1987.tb04387.x

24. Ilarslan H, Dolar FS. Histological and ultrastructural changes in leaves and stems of resistant and susceptible chickpea cultivars to *Ascochyta rabiei*. J Phytopathol. 2002. doi:10.1046/j.1439-0434.2002.00763.x

25. Verma S, Gazara RK, Nizam S, Parween S, Chattopadhyay D, Verma PK. Draft genome sequencing and secretome analysis of fungal phytopathogen *Ascochyta rabiei* provides insight into the necrotrophic effector repertoire. Sci Rep. 2016. doi:10.1038/srep24638

26. Singh K, Nizam S, Sinha M, Verma PK. Comparative transcriptome analysis of the necrotrophic fungus *Ascochyta rabiei* during oxidative stress: Insight for fungal survival in the host plant. PLoS One. 2012. doi:10.1371/journal.pone.0033128

27. Nizam S, Singh K, Verma PK. Expression of the fluorescent proteins DsRed and EGFP to visualize early events of colonization of the chickpea blight fungus *Ascochyta rabiei*. Curr Genet. 2010;56: 391–399. doi:10.1007/s00294-010-0305-3

28. Tsujita K, Suetsugu S, Sasaki N, Furutani M, Oikawa T, Takenawa T. Coordination between the actin cytoskeleton and membrane deformation by a novel membrane tubulation domain of PCH proteins is involved in endocytosis. J Cell Biol. 2006. doi:10.1083/jcb.200508091

29. Kumar V, Fricke R, Bhar D, Reddy-Alla S, Krishnan KS, Bogdan S, et al. Syndapin promotes formation of a postsynaptic membrane system in *Drosophila*. Mol Biol Cell. 2009. doi:10.1091/mbc.E08-10-1072

30. Lanver D, Mendoza-Mendoza A, Brachmann A, Kahmann R. Sho1 and Msb2-related proteins regulate appressorium development in the smut fungus *Ustilago maydis*. Plant Cell. 2010. doi:10.1105/tpc.109.073734

31. Qualmann B, Koch D, Kessels MM. Let’s go bananas: Revisiting the endocytic BAR code. EMBO Journal. 2011. doi:10.1038/emboj.2011.266

32. Fischer-Parton S, Parton RM, Hickey PC, Dijksterhuis J, Atkinson HA, Read ND. Confocal microscopy of FM4-64 as a tool for analysing endocytosis and vesicle trafficking in living fungal hyphae. J Microsc. 2000;198: 246–259. doi:10.1046/j.1365-2818.2000.00708.x

33. Zeigerer A, Gilleron J, Bogorad RL, Marsico G, Nonaka H, Seifert S, et al. Rab5 is necessary for the biogenesis of the endolysosomal system in vivo. Nature. 2012. doi:10.1038/nature11133

34. Steinberg G. Endocytosis and early endosome motility in filamentous fungi. Curr Opin Microbiol. 2014;20: 10–18. doi:10.1016/j.mib.2014.04.001

35. Almeida-Souza L, Frank RAW, García-Nafría J, Colussi A, Gunawardana N, Johnson CM, et al. A Flat BAR Protein Promotes Actin Polymerization at the Base of Clathrin-Coated Pits. Cell. 2018. doi:10.1016/j.cell.2018.05.020

36. Berepiki A, Lichius A, Read ND. Actin organization and dynamics in filamentous fungi. Nat Rev Microbiol. 2011;9: 876–887. doi:10.1038/nrmicro2666

37. Riedl J, Crevenna AH, Kessenbrock K, Yu JH, Neukirchen D, Bista M, et al. Lifeact: A versatile marker to visualize F-actin. Nat Methods. 2008. doi:10.1038/nmeth.1220

38. Park HS, Lee SC, Cardenas ME, Heitman J. Calcium-Calmodulin-Calcineurin Signaling: A Globally Conserved Virulence Cascade in Eukaryotic Microbial Pathogens. Cell Host Microbe. 2019;26: 453–462. doi:10.1016/j.chom.2019.08.004

39. Frost A, Unger VM, De Camilli P. The BAR Domain Superfamily: Membrane-Molding Macromolecules. Cell. 2009. doi:10.1016/j.cell.2009.04.010

40. Frolov VA, Shnyrova A V., Zimmerberg J. Lipid polymorphisms and membrane shape. Cold Spring Harb Perspect Biol. 2011. doi:10.1101/cshperspect.a004747

41. Li X, Gao C, Li L, Liu M, Yin Z, Zhang H, et al. MoEnd3 regulates appressorium formation and virulence through mediating endocytosis in rice blast fungus *Magnaporthe oryzae*. PLoS Pathog. 2017. doi:10.1371/journal.ppat.1006449

42. Leibfried A, Fricke R, Morgan MJ, Bogdan S, Bellaiche Y. Drosophila Cip4 and WASp Define a Branch of the Cdc42-Par6-aPKC Pathway Regulating E-Cadherin Endocytosis. Curr Biol. 2008. doi:10.1016/j.cub.2008.09.063

43. Grove SN, Bracker CE. Protoplasmic organization of hyphal tips among fungi: vesicles and Spitzenkörper. J Bacteriol. 1970. doi:10.1128/jb.104.2.989-1009.1970

44. Li YB, Xu R, Liu C, Shen N, Han LB, Tang D. *Magnaporthe oryzae* fimbrin organizes actin networks in the hyphal tip during polar growth and pathogenesis. PLoS Pathog. 2020;16: 1–27. doi:10.1371/journal.ppat.1008437

45. Peñalva MÁ. Endocytosis in filamentous fungi: Cinderella gets her reward. Curr Opin Microbiol. 2010;13: 684–692. doi:10.1016/j.mib.2010.09.005

46. de Castro PA, Colabardini AC, Manfiolli AO, Chiaratto J, Silva LP, Mattos EC, et al. *Aspergillus fumigatus* calcium-responsive transcription factors regulate cell wall architecture promoting stress tolerance, virulence and caspofungin resistance. PLoS Genetics. 2019. doi:10.1371/journal.pgen.1008551

47. Muñoz A, Bertuzzi M, Bettgenhaeuser J, Iakobachvili N, Bignell EM, Read ND. Different stress-induced calcium signatures are reported by aequorin-mediated calcium measurements in living cells of *Aspergillus fumigates*. PLoS One. 2015. doi:10.1371/journal.pone.0138008

48. Harris SD. Septum formation in *Aspergillus nidulans*. Curr Opin Microbiol. 2001;4: 736– 739. doi:10.1016/S1369-5274(01)00276-4

49. Delgado-Álvarez DL, Bartnicki-García S, Seiler S, Mouriño-Pérez RR. Septum development in *Neurospora crassa*: The septal actomyosin tangle. PLoS One. 2014;9: 29– 30. doi:10.1371/journal.pone.0096744

50. Guertin DA, Trautmann S, McCollum D. (CUL-ID:2436029) Cytokinesis in Eukaryotes. Microbiol Mol Biol Rev. 2002. doi:10.1128/MMBR.66.2.155-178.2002

51. Wu JQ, Kuhn JR, Kovar DR, Pollard TD. Spatial and temporal pathway for assembly and constriction of the contractile ring in fission yeast cytokinesis. Dev Cell. 2003. doi:10.1016/S1534-5807(03)00324-1

52. Martín-García R, Arribas V, Coll PM, Pinar M, Viana RA, Rincón SA, et al. Paxillin-Mediated Recruitment of Calcineurin to the Contractile Ring Is Required for the Correct Progression of Cytokinesis in Fission Yeast. Cell Rep. 2018. doi:10.1016/j.celrep.2018.09.062

53. Lamb CJ, Lawton MA, Dron M, Dixon RA. Signals and transduction mechanisms for activation of plant defenses against microbial attack. Cell. 1989. doi:10.1016/0092-8674(89)90894-5

54. Quest AFG, Bloomenthal J, Bardes ESG, Bell RM. The regulatory domain of protein kinase C coordinates four atoms of zinc. J Biol Chem. 1992.

55. Ichinomiya M, Uchida H, Koshi Y, Ohta A, Horiuchi H. A protein kinase C-encoding gene, pkcA, is essential to the viability of the filamentous fungus *Aspergillus nidulans*. Biosci Biotechnol Biochem. 2007. doi:10.1271/bbb.70409

56. Dichtl K, Samantaray S, Wagener J. Cell wall integrity signalling in human pathogenic fungi. Cell Microbiol. 2016. doi:10.1111/cmi.12612

57. Hurley JH, Newton AC, Parker PJ, Blumberg PM, Nishizuka Y. Taxonomy and function of C1 protein kinase C homology domains. Protein Sci. 2008. doi:10.1002/pro.5560060228

58. Bittova L, Stahelin R V., Cho W. Roles of Ionic Residues of the C1 Domain in Protein Kinase C-α Activation and the Origin of Phosphatidylserine Specificity. J Biol Chem. 2001. doi:10.1074/jbc.M008491200

59. Simunovic M, Manneville JB, Renard HF, Evergren E, Raghunathan K, Bhatia D, et al. Friction Mediates Scission of Tubular Membranes Scaffolded by BAR Proteins. Cell. 2017. doi:10.1016/j.cell.2017.05.047

60. Livak KJ, Schmittgen TD. Analysis of relative gene expression data using real-time quantitative PCR and the 2^-ΔΔCT^ method. Methods. 2001. doi:10.1006/meth.2001.1262

61. Talbot NJ, Ebbole DJ, Hamer JE. Identification and characterization of MPG1, a gene involved in pathogenicity from the rice blast fungus *Magnaporthe grisea*. Plant Cell. 1993. doi:10.1105/tpc.5.11.1575

62. J. Sambrooki D. W. Russell. Molecular cloning : a laboratory manual, III. Red. New York: Cold Spring Harbor Laboratory Press. 2001. doi:10.3724/SP.J.1141.2012.01075

63. Robertson AS, Smythe E, Ayscough KR. Functions of actin in endocytosis. Cell Mol Life Sci. 2009;66: 2049–2065. doi:10.1007/s00018-009-0001-y

